# The dorsal/ventral subdivision of the hindbrain predates the tunicate/vertebrate split

**DOI:** 10.1101/2025.07.15.664975

**Authors:** Matthew J. Kourakis, Yishen Miao, Erin D. Newman-Smith, Kerrianne Ryan, William C. Smith

**Affiliations:** Neuroscience Research Institute, University of California Santa Barbara; Santa Barbara, CA, USA 93106; Department of Molecular, Cellular and Developmental Biology, University of California, Santa Barbara; Santa Barbara, CA, USA 93106; Life Sciences Centre, Dalhousie University; Halifax, NS, Canada B3H 1A5

## Abstract

While the CNSs of the chordate sister clades Tunicata and Vertebrata have unmistakable homology, the extent of their conservation is still being uncovered. We report that the hindbrain of the tunicate *Ciona*, like those of vertebrates, has functionally-distinct dorsal and ventral domains. The *Ciona* dorsal hindbrain functions as a relay and processing center for peripheral sensory neurons, while in vertebrates the dorsal hindbrain forms the cerebellum. Despite the different fates of dorsal hindbrains in *Ciona* versus vertebrates, we present evidence from analysis of gene expression, developmental mechanisms, and neural circuit architecture that they share a common origin. While it is generally agreed that the cerebellum evolved in vertebrates, our findings indicate that a dorsally-positioned precursor to the cerebellum has much deeper roots.

## Introduction

Investigation of vertebrates and their nearest relatives, the invertebrate chordates (*i.e.*, tunicates and cephalochordates), has identified conserved features in their respective central nervous systems that presumably date back to the common chordate ancestor. Evidence for CNS homology between long diverged chordate species takes many forms, including shared anatomy, development, function, neural circuitry, and gene expression. The examination of homologous characteristics helps to delineate the scope and pace of morphological change over time, and to identify the origins of morphological novelty within our own evolutionary lineage. Primitive vertebrate species (*e.g.*, lamprey and hagfish), as well as invertebrate chordates, have been used widely in comparative studies with more complex vertebrates (*1*, *2*). The tadpole larva of *Ciona* is the most widely studied of the invertebrate chordate models, and is the only chordate with a published connectome (*3*). The fully-developed *Ciona* larva, at ∼1 mm in length, is small relative to larvae of amphibians and fish, having only about 2,600 cells, of which approximately 180 comprise the central nervous system (*4*). Despite its diminutive size, the *Ciona* larval CNS drives a complex set of behaviors (*5–7*). The accumulation of data on the *Ciona* larval CNS points to a relatively straightforward correspondence with major CNS subdomains in vertebrates [Fig. 1A; (*8*, *9*)]. In recent decades, much of the investigation of chordate CNS evolution focused on uncovering homologies based on gene expression. For the *Ciona* larval CNS, most expression patterns are supportive of the homologies shown in Fig. 1A, although the anomalous expression of several genes lead to alternative interpretations. For example, the anterior domain of the *Ciona* larval CNS, called the sensory vesicle (SV; also known as the brain vesicle) (Fig. 1a), is divided into anterior and posterior regions (aSV and pSV) that arise from different 8-cell stage lineages (*8*). The expression of a number of genes, including Otx, DMRT, and Lhx2 in the aSV, and Pax3/7, FoxB and Otx in the pSV, point to the aSV being homologous to the forebrain, and the pSV being homologous to the midbrain [reviewed in (*10*)]. However, the absence of Dmbx expression in the pSV, a gene important for vertebrate midbrain development, brought into question the presence of a midbrain homolog in *Ciona* (*11*). However, the accumulation of evidence, supports homology between the pSV and the midbrain including connectomic data which show that the pSV uniquely receives, processes and integrates multimodal sensory inputs (*5*, *10*, *12*, *13*), thus paralleling the function to the vertebrate midbrain tectum.

**Fig. 1.**
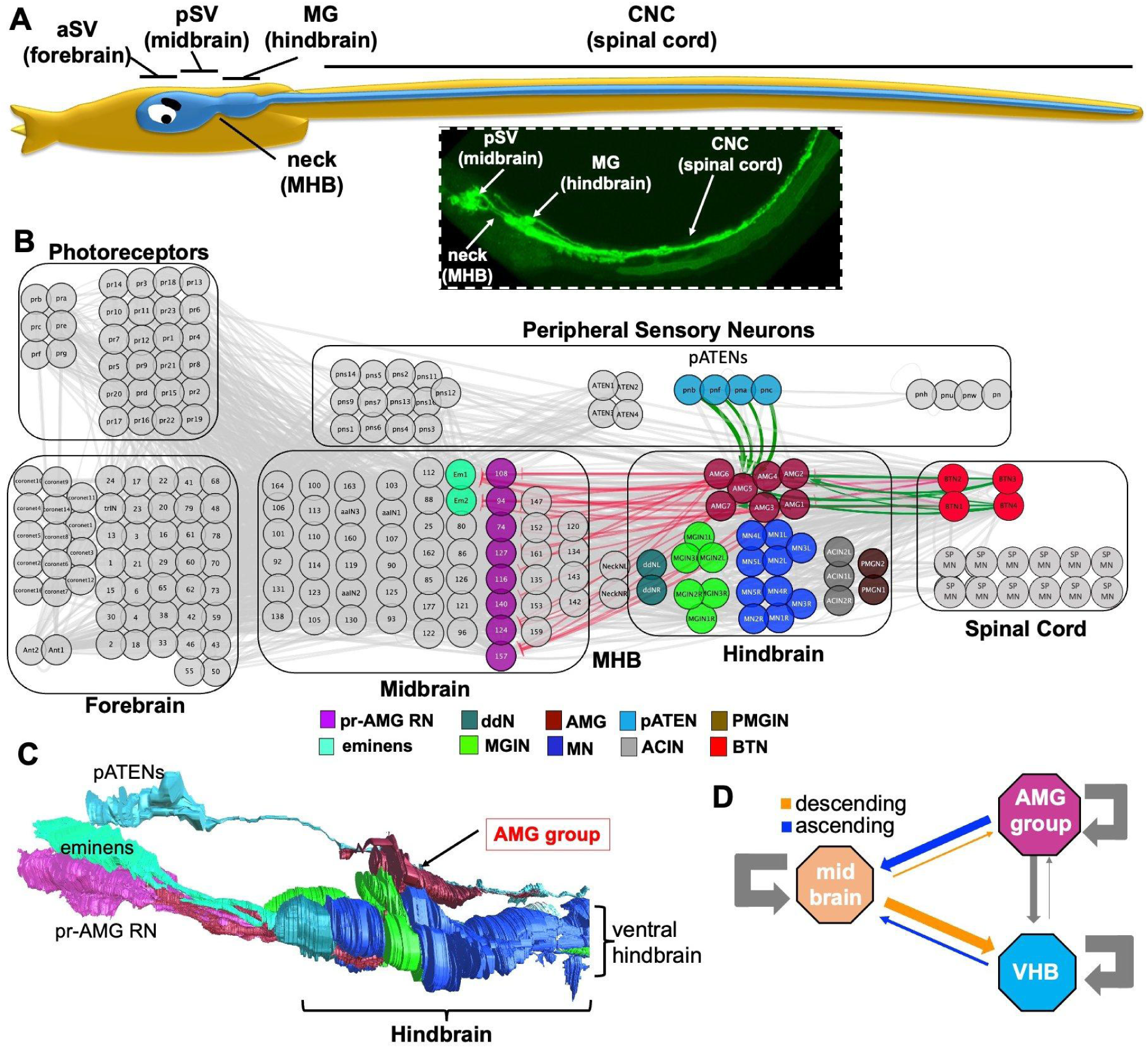
The *Ciona* larval hindbrain has distinct dorsal and ventral domains. **A.** Cartoon of a *Ciona* larva highlighting the central nervous system (blue). CNS domains are labeled with their traditional tunicate names and their putative vertebrate homologs (parentheses). Inset shows major larval CNS subdivisions (excluding the forebrain) labeled with GFP driven by the *cis*-regualtory region of the VACHT gene. **B**. The entire *Ciona* larval CNS connectome highlighting the neurons of the hindbrain, as well as their primary presynaptic inputs (pATENs and BTNs), and postsynaptic targets (eminens and pr-AMG RNs). Green lines indicate putative excitatory synapses, and red lines indicate putative inhibitory synapses. Color coding of neurons is according to (*4*). **C**. Lateral view of the hindbrain reconstructed from serial-section electron micrographs. Also shown are primary inputs to the AMGs, the pATENs, as well as the targets of the AMGs, the pr-AMG RNs and eminens cells. **D**. Summary of the synaptic connectivity between the AMG group, the ventral hindbrain (VHB) and the midbrain. The strengths of all synaptic connections between neurons in the brain regions were summed and then normalized by the number of neurons in each region. These values are reflected in the width of the arrows between the brain regions. Abbreviations: aSV = anterior sensory vesicle; pSV = posterior sensory vesicle; MHB = midbrain-hindbrain boundary; MG = motor ganglion; CNC = caudal nerve cord; pr-AMG RN = photoreceptor-ascending motor ganglion relay neuron; ddN = descending decussating neuron; MGIN = motor ganglion interneuron; AMG = Ascending MG interneuron; MN = motor neuron; pATEN = posterior Apical Trunk Epidermal Neuron; ACIN = Ascending contralateral inhibitory neuron; PMGIN = posterior MG interneuron; BTN = bipolar tail neuron.

Posterior to the pSV is an anatomical constriction in the neural tube called the “neck” (Fig. 1A) which, based on anatomy, gene expression and function, is thought to share homology to the vertebrate midbrain-hindbrain boundary (MHB) (*11*, *14*). Further posterior in the *Ciona* larval CNS are two domains: the first is a thickening in the neural tube called the *motor ganglion* (MG) (also called the “visceral ganglion”), and the second is a narrow and elongated domain extending posteriorly from the motor ganglion, running the length of the tail, called the *caudal nerve cord* (CNC) (Fig. 1a). By anatomy alone, these two domains would appear to correspond to the vertebrate hindbrain and spinal cord, respectively. However, several observations brought into question their anatomical homology to vertebrate structures. For example, it was only recently demonstrated that the *Ciona* CNC contains motor neurons (*9*). The previously perceived lack of motor neurons in the CNC had led to speculation that the MG instead corresponds to the spinal cord - despite the obvious morphological incongruence (*15*). In addition, *Ciona* lacks a member of the Gbx family homeobox transcription factors (*16*), which in vertebrates play critical roles in the development of the MHB and anterior hindbrain, leading to the proposal that *Ciona* larvae lack a hindbrain (*17*, *18*). However, and similar to the pSV, the accumulated evidence of homology, which includes the neuron types of the MG and their synaptic connectivity, agree with hindbrain homology (*9*, *19*). Neurons in the MG include an anterior pair called the *descending decussating neurons* (ddNs), which have circuit homology to the hindbrain Mauthner cells of fish and amphibians (*19*). Additionally, a prominent component of the MG are the six *motor ganglion interneurons* (MGINs), which receive descending motor commands from the pSV, and in turn project to motor neurons in the MG and CNC (*3*, *9*). The connectivity of the MGINs, as well as their expression of the gene Vsx (*20*), suggest homology to vertebrate hindbrain reticulospinal neurons (RSNs). Thus, the anatomy, function, neuron class, synaptic connectivity, and to a varying degree, gene expression, support homology relationships of the *Ciona* MG and CNC to the vertebrate hindbrain and spinal cord, respectively.

We report here further analysis of the *Ciona* motor ganglion, and find that, like the vertebrate hindbrain, it is composed of distinct dorsal and ventral domains which differ in neuron types, gene expression, and function. In vertebrates, this dorsal domain is the cerebellum. In *Ciona*, the dorsal domain is not a cerebellum, but rather a discrete cluster of neurons called the *ascending motor ganglion interneurons* (AMGs). Aside from their shared positional homology (*i.e.*, both structures are found in the dorsal hindbrain), we report that the AMG group shares additional features with the cerebellum, including a dependence on FGF signalling during development, gene expression, and core circuit architecture. Taken together, these observations suggest that the subdivision of the hindbrain into functional dorsal and ventral domains preceded the split of the tunicates from the vertebrates. In tunicates the dorsal domain forms a processing center of peripheral mechanosensory input that bears resemblance to cerebellum-like structures found in various vertebrates.

**Note:** As we have done in recent publications (*12*, *21*), henceforth in this manuscript we will use the names of vertebrate homologs of the *Ciona* CNS domains when referring to them [*i.e.*, hindbrain rather than motor ganglion (MG)]. Fig. 1A shows the correspondences.

## Results

### The Ciona larval hindbrain has distinct dorsal and ventral domains

The *Ciona* larval connectome project identified a total of 30 neurons of six different types in the hindbrain [color coded in Fig. 1b according to (Ryan and Meinertzhagen, 2019)]. Figure 1B shows the *Ciona* larval hindbrain in the context of the entire connectome in which all neurons of the hindbrain are highlighted (colorized). Also labeled are four peripheral sensory neurons called the *posterior Apical Trunk Epidermal Neurons* (pATENs), which are presynaptic to the AMGs (green arrows), and a second major class of input neurons to the AMGs, the bipolar tail neurons (BTNs). The primary targets of the AMGs are in the midbrain, and consist almost exclusively of the eight *photoreceptor-ascending MG neuron relay neurons* (pr-AMG RNs) and the two *eminens* neurons. A reconstruction of the *Ciona* larval hindbrain from the serial EM sections of the connectome project shows that the *Ciona* larval hindbrain is composed of anatomically distinct ventral and dorsal domains, the latter being made up exclusively of the AMG neurons (called henceforth *the AMG group*) (Fig. 1C). The AMG group consists of six inhibitory/VGAT-positive interneurons (AMGs 1,2,3,4,6 and 7), surrounding a central excitatory/VACHT-positive neuron (AMG5) (*13*, *22*)). Functionally, the AMG group acts to receive and relay sensory input from the pATENs to the midbrain pr-AMG RNs and eminens cells [Figure 1B; (*22*)]. In contrast to the dorsal domain, the ventral hindbrain domain receives and executes sensorimotor commands from descending excitatory and inhibitory relay interneurons of the midbrain (*13*). The primary midbrain synaptic targets in the hindbrain are the RSN-like MGINs, which in turn project to motor neurons in the hindbrain and spinal cord (*3*, *9*). Also in the *Ciona* larval ventral hindbrain are the ddNs, mentioned in the Introduction, as well as the *ascending contralateral inhibitory neurons* (ACINs), and the *two posterior MG interneurons* (PMGINs). The ACINs are thought to be components of the swimming central pattern generator (*23*), while the function(s) of the PMGINs is unknown. The distinction between functions of the AMG group and the ventral hindbrain is particularly evident when their connectivities to each other, and to the midbrain, are compared (Fig. 1D). While the ventral hindbrain receives extensive descending input from the midbrain, the AMG group receives very little, and is instead characterized by its extensive ascending projections to the midbrain, consistent with its role as a relay center for peripheral sensory inputs. While there is extensive synaptic connectivity within the AMG group and the ventral hindbrain (gray arrows in Fig. 1D), there is very little between the two, and the few synaptic connections that are present consist almost exclusively of projections from the AMG group to the ventral hindbrain.

### The AMG group processes mechanosensory input

Epidermal sensory neurons (*e.g.*, the pATENs) in *Ciona* larvae are thought to be either mechanosensitive, chemosensitive, or both (*22*). We observe that touching the pATENs of a larva with a fine probe results in an evoked swim, confirming that they are mechanosensory, although they are also reported to be chemosensory (*24*) (Figure 2A; three additional examples are shown in Movie S1). The four pATENs are glutamatergic (*13*), and project almost exclusively to the sole excitatory AMG, AMG5 [Figure 2B; (*22*)] (*13*). AMG5 has a large bifurcating axon (Figure 2C) not found in the other AMGs [see *Source data 1, Figure 1* of (*3*) for serial-section EM reconstructions of all larval neurons], and is presynaptic to the inhibitory AMGs, with the apparent exception of AMG2 [Figure 2B; (*3*)]. Output from the AMG group to the midbrain is via the ascending projections of the inhibitory AMGs (Figure 2B). Using a Hox10 *cis*-reglatory region driving GFP (*25*) the ascending axons of AMG2 and 3 are evident (Figure 2D, yellow arrows). Thus, the AMG group changes the valence of the mechanosensory input of the pATENs from excitatory (glutamate via acetylcholine) to outgoing inhibitory (GABA).

**Fig. 2.**
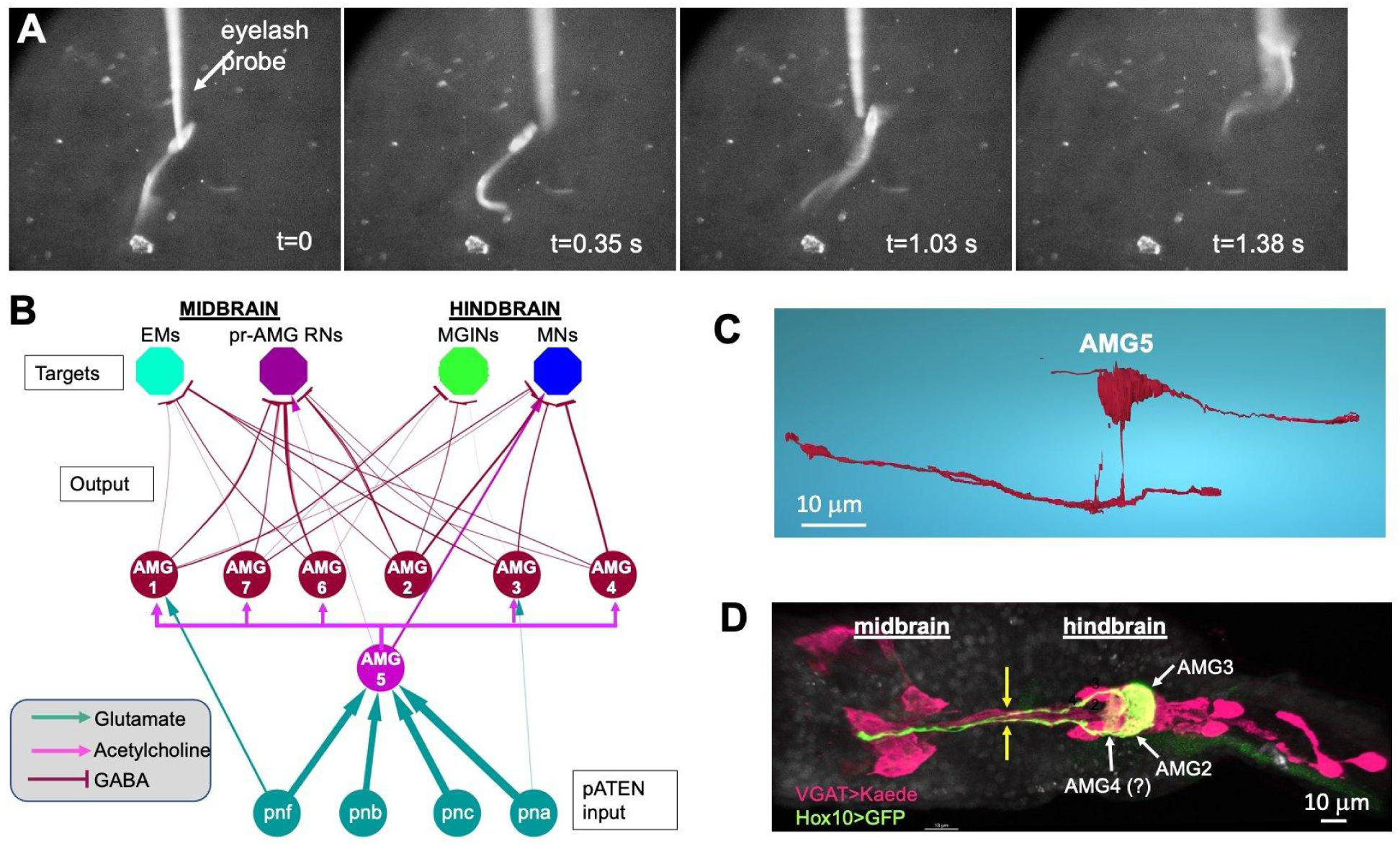
Activity and circuitry of the AMG group. **A**. Response of a *Ciona* larva to mechanical stimulation (*i.e.*, touch) of the pATENs. s= seconds. **B**. Neural circuit of the AMG group. Input from the mechanosensory pATENs (*pnf*, *pnb*, *pnc* and *pna*) primarily target AMG5 (cholinergic), which then branches to inhibitory output AMG neurons. **C**. Reconstruction of AMG5 from serial-section electron microscopy. **D.** Ascending projections from AMGs 2 and 3 labeled with a Hox10 promoter>GFP plasmid (green). Also shown are VGAT-positive neurons (magenta). Faint green fluorescence observed anterior to AMG2 and 3 may be AMG 4. (see Fig. 1 legend for abbreviations). Dorsal view is shown, anterior is left.

In addition to receiving direct mechanosensory input from the pATENs at AMG5, the AMGs are postsynaptic to the four *bipolar tail neurons* (BTNs), which function as relay interneurons for the mechanosensory *dorsal caudal epidermal neurons* (DCENs), which are found along the tail (Figure 3A). The BTN neurons are thought to be homologous to neurons of the vertebrate dorsal root ganglia (*2*), consistent with their hypothesized role of providing proprioceptive input on tail movements (*26*, *27*). While previous studies have reported the presence of a single anterior GABAergic BTN and single posterior cholinergic BTN (*28*), we find two anterior VGAT-positive (VGAT^+^) BTNs (BTN1 and 2), and two posterior VACHT^+^ BTNs (BTN3 and and 4) (Fig. 3B). Thus, the AMG group appears to receive both positive and negative representations of tail movements. The potential significance of BTN input for the signal processing functions of the AMG group, and how this may tie the AMG group to cerebellum-like structures found in certain vertebrates, is presented in the Discussion.

**Fig. 3.**
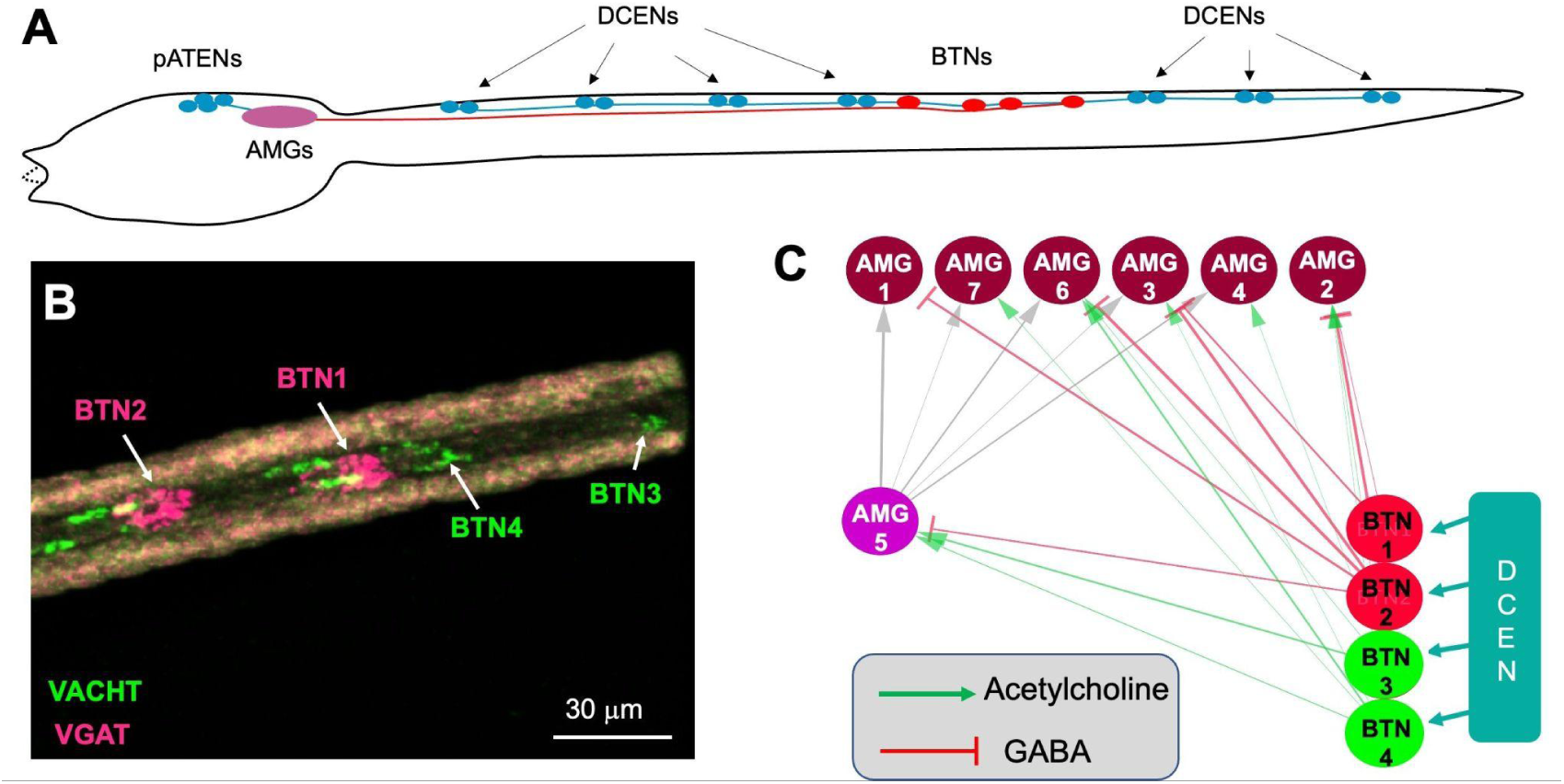
Bipolar tail neurons and the AMG group. **A.** Relative positions of the pATENs, DCENs, AMGs and BTNs in a *Ciona* larva [after (*22*)]. **B.** BTNs labeled by in situ hybridization for VACHT (green) and VGAT (magenta). **C.** Synaptic input from the BTNs to the AMG group. See Fig 1. legend for abbreviations.

### Gene Expression in the AMG group

The above sections detail that the dorsal and ventral hindbrain of *Ciona* larvae can be distinguished by anatomy, cell-type composition, function and circuit architecture. These differences will, of course, be reflected in differential gene expression between the two domains. In vertebrates, the dorsal domain is the cerebellum. Recognizing the apparent similarity in the partitioning of the hindbrain in both vertebrates and *Ciona* to dorsal and ventral domains, we started our investigation of the AMG group by examining genes expressed in the cerebellum, starting with the transcription factors Lhx1/5 and Otp, which are expressed in non-overlapping subpopulations of inhibitory neurons in the cerebellum (*29*). We find that in *Ciona* larvae both Lhx1/5 and Otp are expressed in the inhibitory AMGs, with Lhx1/5 expressed in the anterior two AMGs (AMG6 and 7) (Fig. 4A), and Otp expressed posteriorly in AMGs 2,3 and 4 (Fig. 4B).

**Fig. 4.**
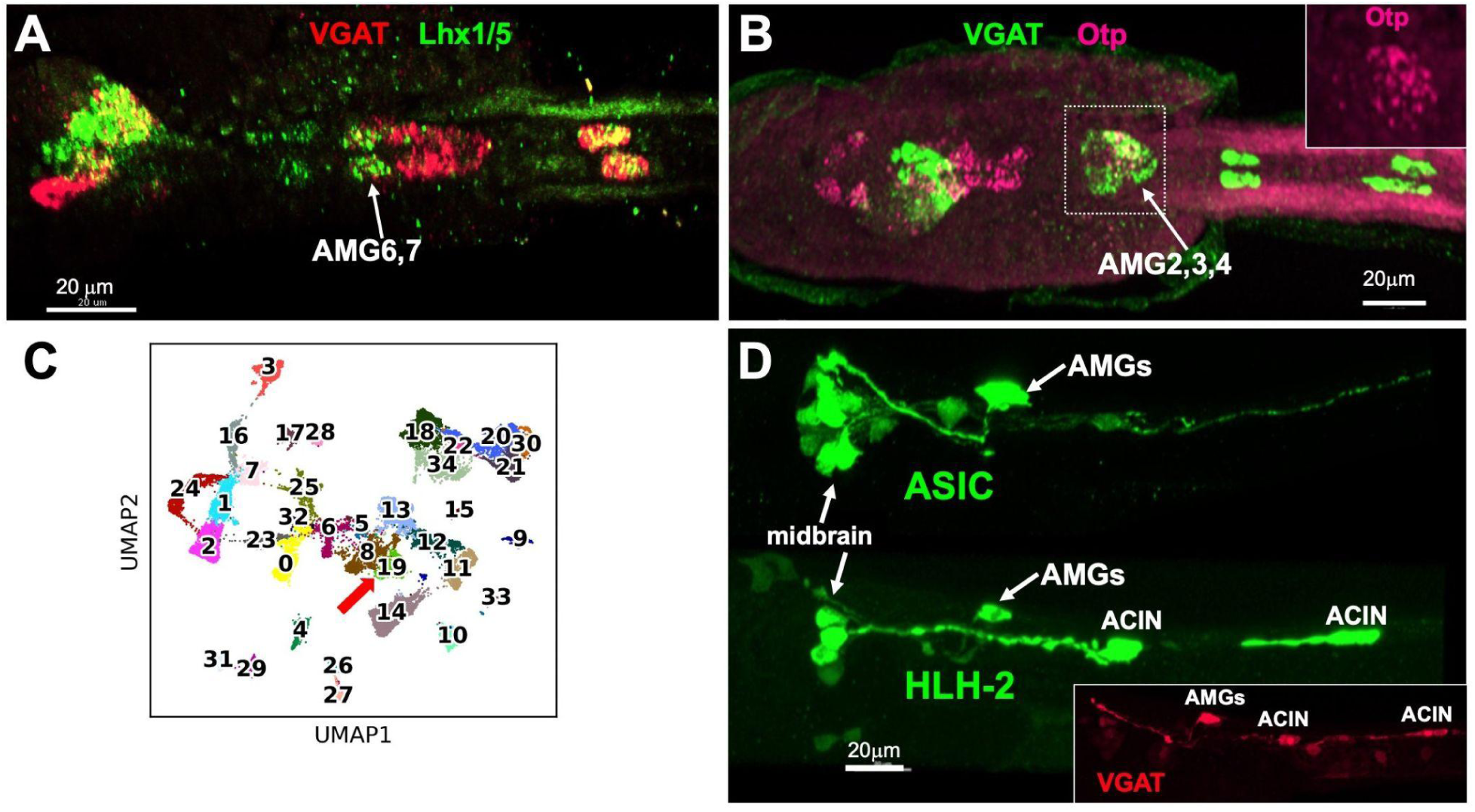
Gene expression in the AMG group. **A and B**. Expression of Lhx1/5 in a *Ciona* larva by in situ hybridization (panel A, green), and Otp (panel B; magenta). Both panels also show expression of VGAT. **C**. Cluster analysis of single-cell transcriptomes from combined datasets of *Ciona* larvae (*30*, *31*). The red arrow indicates the candidate cluster corresponding to the inhibitory AMGs. **D.** Larval stage expression of ASIC and NSCL-2 GFP reporter constructs. **A** and **B** are dorsal views, and **D** is lateral.

To gain a fuller picture of the transcription profile of the AMGs, we took advantage of the availability of two single-cell RNAseq datasets for *Ciona* larvae (*30*, *31*). As detailed in the Material and Methods, these two datasets were combined to make a single dataset. Differential gene expression analysis identified a single cluster coexpressing VGAT, Otp, and Lhx1/5 (cluster 19 in Fig. 4C; Supplemental Table 1), suggesting that it may correspond to the inhibitory AMGs. The presence of differentially expressed Otp and Lhx1/5 in the same cluster suggests that the Otp^+^ and Lhx1/5^+^ AMGs are very similar. To validate cluster 19 as corresponding to the inhibitory AMGs, we examined the expression of two of its most highly differentially expressed genes: the *Ciona* orthologs of the *Acid-sensing ion channel* (ASIC) (*32*), and *Helix-loop-helix protein 2* (HLH-2) (STable 1). To assess the expression of ASIC in *Ciona* larvae we used a previously described expression construct with the *CiASICb* enhancer driving GFP (*32*), while for HLH-2 we made a new construct with ∼1.5 kb 5’-flanking region driving GFP (see Material and Methods). We observed strong expression from both the ASIC and HLH-2 expression constructs in the AMGs, as well as elsewhere in the CNS. We note, however, that the sites of expression outside the AMGs differ between the two constructs, as might be expected for genes identified specifically for their differential expression in the AMGs. The HLH-2 construct also drove expression in the ACINs of the ventral hindbrain, which can be identified by their posterior placement within the ventral hindbrain and their expression of VGAT (Fig. 4D insert). These expression patterns are consistent with cluster 19 corresponding to the inhibitory AMGs. In addition, both ASIC and HLH-2 are differentially expressed in Purkinje (*i.e.*, inhibitory) neurons of the mammalian cerebellum [STable 1; (*33–36*)]. Finally, we lack sufficient expression data on AMG5, other than it expresses VACHT (*13*), to perform a similar analysis. Moreover, because there is only one AMG5 neuron per larva, single-cell transcriptome analysis may not yield sufficient data to allow for its identification.

### Development of the AMGs

We investigated the timing of the inhibitory AMG neuron development by examining the onset of expression of VGAT, determined by in situ hybridization. At mid-tailbud stages (stage 23) no expression of VGAT was observed in the developing hindbrain, although expression was detected in the developing inhibitory BTNs of the tail. However, at this stage VACHT was detected in the hindbrain (Fig. 5A). Because there is only one VACHT^+^ neuron in the dorsal hindbrain, most, or all, of the VACHT expression at this stage is in the ventral hindbrain. By stage 24, VGAT expression was detected in the dorsal hindbrain (Fig. 5 B). This pattern of expression, with the VACHT^+^ hindbrain neurons developing before the VGAT^+^ neurons, was observed in each of four samples examined. The timing of these expression patterns, including the earlier detectable VGAT in the caudally positioned BTNs, as well as timing of VGAT/VACHT in the midbrain could give clues as to how functional circuits for sensorimotor pathways are established temporally. The published connectome is for a *Ciona* larva, earlier stage-specific connectomic data could provide insight into the sequential development of neural circuits.

**Fig. 5.**
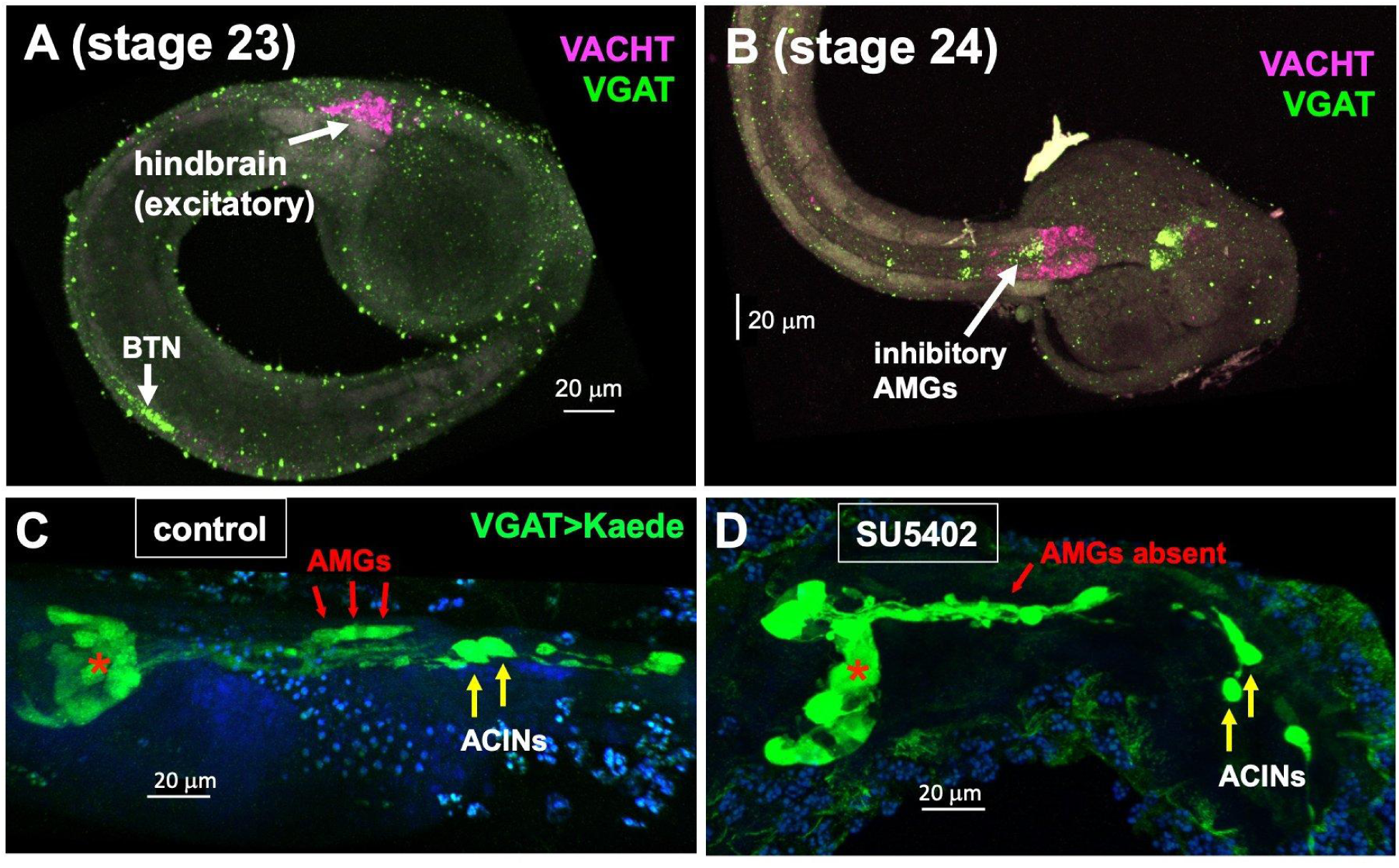
Development of the AMG group. **A and B**. Expression of VGAT (green) and VACHT (magenta) in the developing hindbrain. Anterior is to the right. **C.** Vehicle control for SU5402 experiment (DMSO treatment). Larvae expressing Kaede fluorescent protein driven by the VGAT promoter and labeled with anti-Kaede primary antibody and an AlexaFluor secondary (red arrows). **D**. SU5402 treatment eliminates the VGAT-expressing AMGs. Red asterisks in panels A and B indicate midbrain VGAT^+^ neurons. Abbreviations. BTN = bipolar tail neurons; AMG = *ascending motor ganglion interneurons*. ACIN = ascending contralateral inhibitory neurons. In **C** and **D** anterior is to the left.

In vertebrates, the dorsal hindbrain is induced via the action of FGF8 produced at the MHB (*37*). The *Ciona* ortholog of this gene, called FGF8/17/18, shows a remarkably well-conserved expression pattern in the developing CNS, being expressed in the CNS neck region, immediately posterior to neurons expressing *OTX* and *engrailed (En)*, as in vertebrates (*38*). The naming of FGF8/17/18 reflects the multiple vertebrate orthologs of the single *Ciona* gene, resulting from independent whole-genome duplications that occurred in the vertebrate lineage, after vertebrate and tunicate lineages diverged (*39*). Based on morphology, gene expression, and inductive activity, the *Ciona* larval CNS neck region has been equated with the MHB (*40*, *41*). Previous studies showed that the loss of FGF8/17/18 disrupts motor ganglion gene expression (*14*), although disruptions specific to the AMG group were not addressed. To investigate a possible role for FGF in the development of the AMG group, *Ciona* embryos were treated with the FGF receptor inhibitor SU5402. To assess the presence/absence of the AMG group, experiments were performed in a stable transgenic line expressing Kaede fluorescent protein under the control of the VGAT promoter (*42*), which labels the six inhibitory AMG neurons (1-4, and 6-7). We found that treatment of the embryos for 30 minutes with 20 𝜇mole SU5402 at the late neurula stage eliminated VGAT expression in the dorsal hindbrain, while a vehicle control (DMSO) had no effect (Fig. 5 C and D). Despite the loss of the AMGs, other VGAT-expressing CNS neurons appeared to be present, including those in the midbrain (red asterisks) and the ACINs of the motor ganglion, indicating that the AMGs are uniquely induced at this developmental stage. Treatment of older embryos with SU5402, from the initial tailbud stage onward, had no effect on the development of the AMG group (not shown).

### Inhibitory AMGs have spontaneous spiking activity

Given the similar anatomical location, gene expression, circuit architecture, and development of the AMG group and neurons of the vertebrate dorsal hindbrain/cerebellum, we extended our analysis to determine whether the inhibitory AMG neurons might display spontaneous spiking activity, as has been observed in some inhibitory neurons of the cerebellum, namely the Purkinje cells. In vertebrates this spontaneous activity consists of both high frequency “simple spikes” (>40 Hz), and trains of slower (1-3 Hz) “complex spikes” (*43–45*). In the present experiment, 1-cell stage *Ciona* embryos were electroporated with a plasmid containing GCaMP6f driven by the VGAT promoter (VGAT>GCaMP6f), and then assessed at the larval stage (*12*). The temporal resolving power of calcium transients by GCaMP is below what is needed to detect simple spikes, but we have successfully used GCaMP to quantify rhythmic firing behavior of other neurons in *Ciona* larvae on the time-scale of complex spikes (*12*). Fig. 6A shows two frames from a representative recording of GCaMP activity in which a spike (calcium transient) was observed in the AMG group (arrows) (see Movie S2). Five GCaMP expressing larvae were imaged, and Fig. 6B, shows a representative plot of normalized fluorescence (ΔF/F_0_) versus time. In the five larvae analyzed (fig. S1), we observed trains of calcium transients, sometimes persisting through the imaging sessions (∼3 minutes), but in other cases occurring in bursts lasting tens of seconds (*e.g.*, larva 4 fig. S1). In analyzing fluorescence data we selected small regions of interest (ROI) for quantification that correspond to the approximate size of individual cells. While it is possible these ROIs contained more than one cell, we also observed temporally matched calcium transients in separate AMGs (larva 1 fig. S1), indicating that they are coupled. The mean spiking frequency for the inhibitory AMG neurons from five larvae was 0.42 ± 0.23 Hz (Fig. 6C). Overall, the pattern of calcium transients in the inhibitory AMGs appears similar to complex spikes (*46*), although their frequency is lower. The lower frequency may indicate different mechanisms between the inhibitory AMGs and the Purkinje cells, and that these spontaneous activities may be unrelated. However, it is also well documented that the frequency of rhythmically activity neurons is temperature dependent (*47*), and *Ciona* were imaged at 18° C.

**Fig. 6.**
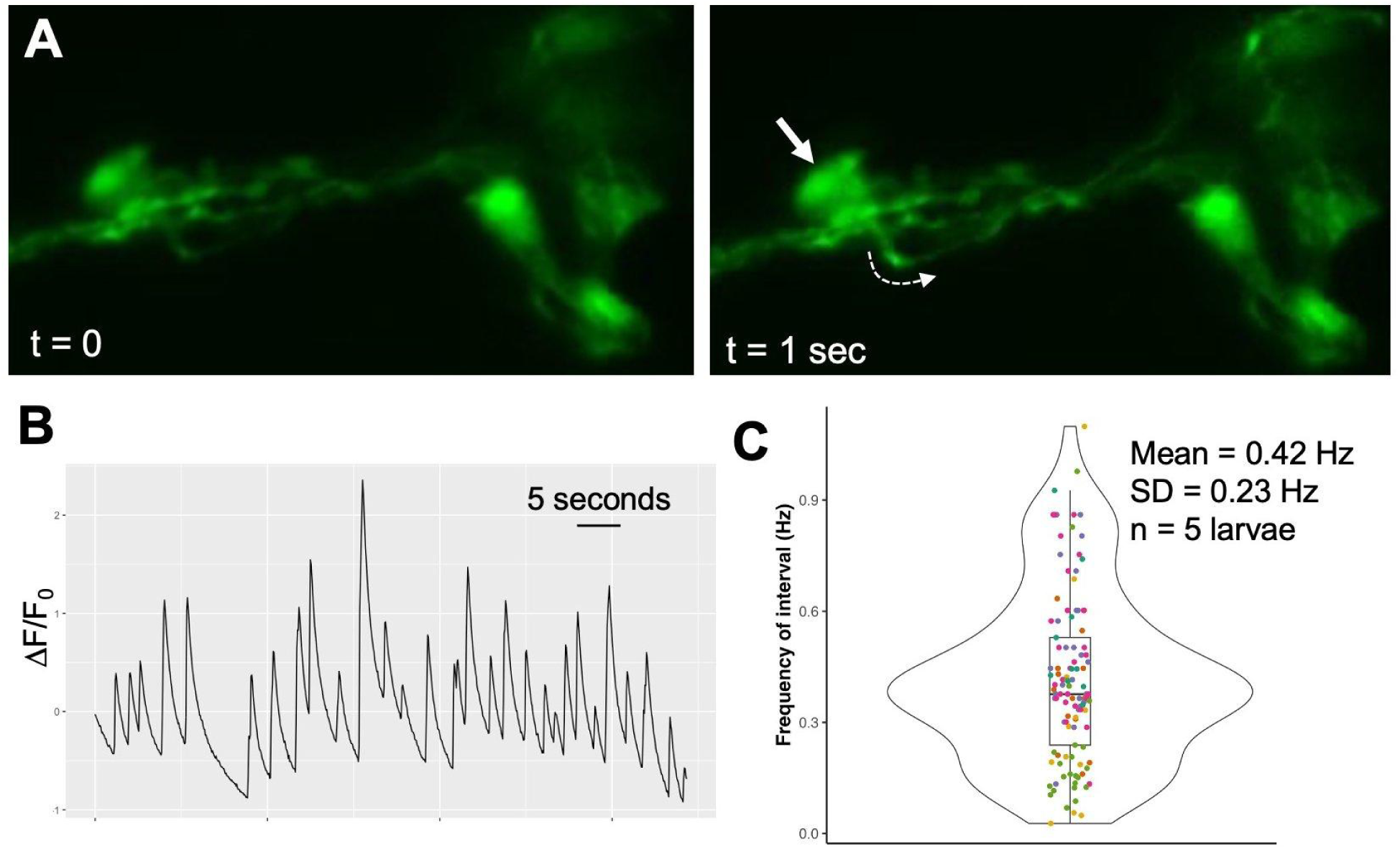
Spontaneous activity in the inhibitory AMGs. **A**. Representative example of a calcium transient in the AMG group detected with GCaMP6f driven by the VGAT promoter. The calcium transient is observed at t=1 sec as increased fluorescence in the AMG group (arrow) and in an ascending axon (dashed arrow). Images are frames taken from Movie S2. Anterior is to the right. **B**. Representative plot of a GCaMP6f fluorescence in the inhibitory AMG group. **C**. Combined data on spontaneous activity in the AMG group from five larvae. Data from each larva are a different color, and dots correspond to all 1/peak interval times in seconds for each recording, and were used to determine mean frequency in Hz. The box indicates the median values (middle bar) and first to third interquartile ranges (upper and lower bars); whiskers indicate 1.5× the interquartile ranges.

## Discussion

### The Ciona larval hindbrain

As detailed in the Introduction, multiple lines of evidence support the homology of the *Ciona* larval MG with the vertebrate hindbrain. These include its anatomical location in the *Ciona* larva between the MHB and the spinal cord, shared gene expression, constituent neuron type, and synaptic connectivity (*8*, *9*, *19*, *20*). Nevertheless, in *Ciona* the developmental expression of several genes appear to be confounding - such as the absence of a Gbx ortholog, and the atypical expression of Hox10 within the MG (*25*). In both vertebrates and cephalochordates (*e.g., amphioxus*) - the outgroup for tunicates and vertebrates (*48*), Hox10 paralogues are expressed in the caudal CNS (*e.g.,* lumbar spinal cord for vertebrates) (*49*, *50*). Thus, it is most likely that *Ciona* Hox10 expression in the hindbrain is derived within the chordates, consistent with the overall disruption, dispersal and modification of the *Ciona* Hox cluster (*25*, *51*). However, in light of the multiple lines of evidence supporting the homology of the MG and the hindbrain, these differences likely point to plasticity and drift of developmental mechanisms, reflecting changes which have accrued separately in the lineages leading to *Ciona* and to vertebrates, rather than to independent origins of the MG and the hindbrain. A similar mix of conservation and divergence is observed, for example, in comparing gene regulatory networks between tunicate and vertebrate notochords (*52*). Moreover, such drift may be expected given the large evolutionary distance between tunicates and vertebrates, and the many apparent gene losses and rearrangements in the *Ciona* genome when compared to those of other chordates (*51*, *53*). In addition, given the conserved gene expression, morphology, function and neuron composition of the CNC to the vertebrate spinal cord (*9*, *54*, *55*), there appears to be little support for the earlier hypothesis that the MG has homology to both the vertebrate hindbrain and spinal cord (*15*). Nevertheless, the *Ciona* hindbrain is a vastly simpler structure than its vertebrate counterpart, having only thirty neurons (Fig. 1B). Accordingly, the *Ciona* larval hindbrain lacks evident subdivisions ventrally, such as pons and medulla oblongata, as found in vertebrates. However, one subdivision that the *Ciona* hindbrain does have is in the dorsal/ventral axis. In *Ciona*, the dorsal and ventral domains differ in function, gene expression, neuron type and synaptic connectivity. Moreover, it is thought that the dorsal and ventral hindbrains of the *Ciona* larvae develop from different embryonic lineages (*56*).

The gene regulatory pathways of motor- and interneuron specification in the ventral *Ciona* larval hindbrain have been extensively characterized (*57–59*). In the dorsal hindbrain, the specification of AMG5 (the sole cholinergic AMG) requires the transcription factor *Ebf* (*56*). We show here that the GABAergic AMG neurons are transcriptionally heterogeneous, with the anterior two (AMG6 and 7) expressing the transcription factor Lhx1/5, the middle three (AMG2,3,4) expressing the transcription factor *Otp*, while the most posterior (AMG1) expresses neither of these genes. Whether this heterogeneity of gene expression in the GABAergic AMGs reflects functional differences is unknown, but at the level of synaptic connectivity (Figs. 2B and 3C) they appear to be equivalent. It is also unknown if either or both Otp and Lhx1/5 are necessary for inhibitory AMG development, although they all appear to be induced by the same FGF-dependent pathway (Fig. 5). Our identification of a scRNAseq cluster for the inhibitory AMGs provides a tool for future investigation of AMG development and function.

### Did dorsal/ventral subdivision of the hindbrain arise independently in vertebrates and tunicates?

Anatomically, the AMG group occupies an equivalent position in the CNS to the cerebellum of vertebrates (*i.e.*, dorsal hindbrain). However, the simple structure of the AMG, and its function as a relay center for mechanosensory input, clearly distinguishes it from a cerebellum. Moreover, the cerebellum is thought to have emerged following the split of jawed vertebrates (gnathostomes) from the jawless vertebrates (agnathans) (*60–62*). Nevertheless, embryos of lamprey and hagfish (both agnathans) express the genes *Ptf1a* and *Atonal* in the developing dorsal hindbrain (rhombomere 1) (*63*). Both of these genes are commonly used markers of GABAergic and glutamatergic neurons on the cerebellum, respectively, although they both are widely expressed elsewhere in the CNS. While the dorsal rhombomere 1 derivatives appear to degenerate in hagfish, the Ptf1a positive cells of lamprey appear to develop into inhibitory neurons, paralleling certain aspects of cerebellum development.

However, the dorsal hindbrain domain of lamprey has been described as comprising an undifferentiated plate-like cerebellum (*64*). This would appear to agree with single-cell RNA-sequence analysis of the lamprey CNS. While many lamprey neuron types could be tied to mouse neuron types based on transcriptomics, no correspondence of lamprey neurons to cerebellar neurons were found (*61*). Thus, it appears that while the agnathans possess a distinct dorsal hindbrain domain which express molecular markers associated with the cerebellum, this domain may lack other cerebellum-defining characteristics. A connectome for lamprey or hagfish, though currently lacking, could supplement expression and RNAseq data, revealing whether there exist commonalities in circuit logic between dorsal hindbrains of cyclostomes and vertebrates. Despite this, conserved gene expression indicates that the presence of distinct dorsal and ventral hindbrain domains predates the gnathostome/agnathan split, with the dorsal hindbrain of the common gnathostome/agnathan ancestor serving as a possible precursor to the cerebellum.

We hypothesize, based on data presented here, that the appearance of dorsal and ventral hindbrain domains is even more ancient, predating even the vertebrate/tunicate split. The alternative, that the *Ciona* dorsal hindbrain domain (*i.e.*, AMG group) evolved independently from those of vertebrates, appears less plausible given the weight of evidence. In the scenario we propose, the common ancestor already had a dorsal/ventral-regionalized hindbrain; after long and separate evolutionary trajectories, the dorsal domain gave rise to the cerebellum in the lineage leading to vertebrates, and the AMG complex in the tunicates. We find conserved gene expression between the AMG group and the cerebellum. While there are no known genes that are exclusive to the cerebellum, we present here are a set of genes differentially expressed in both, including Lhx1/5, Otp, ASIC, VGAT, and HLH-2 (Figure 4). In addition, cluster 19 contains a number of other genes that are known to be differentially expressed in cerebellum (STable 1, green highlights). One transcription factor that is conspicuously absent in the AMG group is Ptf1a, which is a key regulator of GABAergic neurons in the cerebellum, and elsewhere (*37*, *65*). While *Ciona* was purported to have a Ptf1a ortholog (*66*), further analysis showed that the gene annotated as *Ciona* Ptf1a is a paralog, not an ortholog, of mammalian Ptf1a, and lies within the Fer cluster of the larger Ptf1a-related gene family (*67*). However, the cephalochordate *Branchiostoma lanceolatum* does appear to have a true Ptf1a ortholog (fig. S2). Because the cephalochordate subphylum of the chordates is basal to both the tunicates and vertebrates (*48*), it appears most likely that the tunicates lost the Ptf1a gene. Whether cephalochordates have a distinct dorsal hindbrain domain, and whether it expresses Ptf1a, has not been explored.

We performed a comparison of the transcriptomes of all *Ciona* larval clusters shown in Fig. 4C to a set of mouse cerebellum scRNAseq transcriptomes (*68*). Briefly, we identified one-to-one orthologs between the mouse and *Ciona* genomes, and used these orthologs to map mouse gene expression to corresponding *Ciona* genes, and measured the cosine similarity between mouse and *Ciona* clusters (see Materials and Methods). While there were varying degrees of match between human and *Ciona* single cell transcriptomes, none of the *Ciona* clusters was a good match for the predominant cell types in the core circuit of the cerebellum (Purkinje and granule; fig. S3). While this result appears similar to lamprey, the enormous evolutionary distance between tunicates and vertebrates, as well as the much greater neuron-type diversity of vertebrates compared to tunicates, make this approach of questionable validity.

We conclude that the gene expression data presented here is consistent with the hypothesis that the AMG group has a common origin with the dorsal hindbrains of vertebrates. The absence of definitive markers of cerebellum prevents a stronger conclusion. However, as discussed below, examination of the neural circuitry of the AMG group suggests conserved processing motifs between the AMG group and cerebellum.

### Is the AMG group a cerebellum-like structure?

The *Ciona* larva has a complex peripheral nervous system that projects to the CNS at several relay centers, the AMG group being one of them (*22*). However, the circuitry of the AMG group stands apart, and suggests a circuit architecture similar to those found in the vertebrate cerebellum-like structures, although on a vastly simplified scale, and with some notable differences. Vertebrate cerebellum-like structures are anatomically and functionally distinct from the cerebellum and receive, process, and relay direct sensory inputs from both mechano- and electrosensory neurons (*69–72*). Examples include a cerebellum-like structure found in the dorsal cochlear nucleus of mammals that receives and processes mechanosensory input from the auditory nerve, and a cerebellum-like structure found in the medial octavolateral nucleus of fish and amphibians that receives input from the mechanosensory lateral line (*73*, *74*). The core neural circuits of the cerebellum and cerebellum-like structures are similar, with excitatory granule cells (or granule-like cells in cerebellum-like structures) branching to a network of parallel fibers making multiple synaptic contacts with inhibitory Purkinje/Purkinje-like cells. In the *Ciona* AMG group, inputs from the mechanosensory pATENs project to the central cholinergic neuron, AMG5 (Fig. 2B). AMG5, with its large bifurcating axon, then branches to form synapses with the inhibitory AMGs (Fig. 2B and C). Also as in cerebellums and cerebellum-like structures, the output is inhibitory (e.g., Purkinje cells in the cerebellum and the inhibitory AMGs in *Ciona*). Thus overall, the circuit architecture of the AMG group with a branching excitatory input neuron synapsing to multiple inhibitory output neurons appears similar to cerebellum and cerebellum-like structures, although functionally the AMG group appears to have more in common with cerebellum-like structures. In this circuit, the inhibitory AMG neurons would be functional equivalents of Purkinje cells, while AMG5 would have a function akin to granule cells. Despite this similarity, important differences are evident as well; the most obvious being that in cerebellum and cerebellum-like structures the core processing circuit is repeated on a numerically vast scale, while in the AMG group there is only a single circuit consisting of a single excitatory input neuron and six inhibitory output neurons. However, it should be noted that the small number of neurons in the AMG group is in keeping with the overall small number of neurons in all classes in *Ciona* larvae (*4*), and is perhaps a reflection of a simplification of tunicates from a more complex ancestor (*75*).

A second noticeable difference between the AMG group and cerebellum/cerebellum-like structures is that AMG5 is cholinergic while the granule cells are glutamatergic - although both are excitatory. While this may indicate that AMG5 and granule cells have different origins, it is well documented that homologous neurons can switch neurotransmitter use (*76*). It is also worth noting that *Ciona* larvae are overall lacking glutamatergic interneurons; the only glutamatergic neurons in *Ciona* larvae are sensory (photoreceptors, antennae cells, and peripheral neurons) (*13*). However, glutamatergic interneurons are present in cephalochordates (*amphioxus*), suggesting that the lack of glutamatergic interneurons may be a derived feature of tunicates (*77*), and that for unknown reasons, tunicates have either lost glutamatergic interneurons, or changed their neurotransmitter use. This change from glutamatergic to cholinergic may also be evident in the putative *Ciona* reticulospinal neurons (*9*, *19*), which are cholinergic, rather than glutamatergic as in vertebrates (*13*, *78*).

Functionally, cerebellum-like structures not only receive sensory input, but also act as predictive cancellation filters (*74*, *79*, *80*). Cancellation filters are essential for countering sensory input, known as reafference, that arise from self-stimulation of mechanosensory neurons by the animal’s own motor behavior. The cerebellum-like structures receive corollary discharge and proprioceptive inputs which are then used to predict or subtract self-generated signals (*81–83*). It is likely that *Ciona* larvae would face the same self-stimulation problem as observed in other animals, both invertebrate and vertebrate (*84*). Examination of the *Ciona* connectome dataset reveals a plausible predictive cancellation filter functionality in the AMG group. The AMGs, in addition to receiving direct mechanosensory input from the pATENs at AMG5, are postsynaptic to the four *bipolar tail neurons* (BTNs) (Fig. 3C). The BTNs serve as relay interneurons for the mechanosensory *dorsal caudal epidermal neurons* (DCENs) (*22*), which have a proposed proprioceptive function (*27*). Significantly, the BTNs have mixed valence, with the anterior two (BTN1 and 2) being VGAT^+^ (*i.e.*, inhibitory), while the posterior two (BTN3 and 4) are VACHT^+^ (*i.e.*, excitatory) (Fig. 3B). The output from the BTNs should thus provide both positive and negative representations of tail movements to the AMG group, as is seen in cerebellum-like structures. Whether the BTN input to the AMGs functions in reafference cancellation remains to be determined.

In summary, we present here a new perspective on the evolution of the cerebellum: the *Ciona* larval CNS not only possesses a domain that shares homology to the vertebrate hindbrain (as we outline in the Introduction), but we further specify in this communication that this homology includes functionally distinct dorsal and ventral domains. The features tying the AMG group to the vertebrate dorsal hindbrain are numerous, and include gene expression, developmental induction, circuit architecture and spiking activity. Taken as a whole, these data suggest that the cerebellum evolved from a structure that was already present before the tunicate/vertebrate split. Moreover, it has been proposed that the cerebellum evolved from a cerebellum-like structure (*70*, *72*), suggesting that the cerebellum evolved from a structure that was already partially prefigured. Connectome data on the hindbrains of both lamprey and cephalochordates would likely be informative. In lamprey, we predict that the circuit architecture of the dorsal hindbrain would resemble a cerebellum-like structure. Connectome data from cephalochordates could indicate if the dorsal and ventral hindbrain structures were present prior to the split of the cephalochordates from the olfactors (tunicates and vertebrates).

## Material and methods

### Animals

Adult *Ciona robusta* (aka, *C. intestinalis* type A) were collected at the Santa Barbara Harbor with the exception of the stable VGAT>Kaede line (Tg[MiCiVGATK]2, which was obtained from CITRES https://marinebio.nbrp.jp/ciona/) and cultured at the UC Santa Barbara Marine Laboratory. Larvae were generated by dissecting gametes from adults. VGAT>Kaede transgenic larvae were produced by crossing wild type eggs with frozen VGAT>Kaede sperm aliquots.

### HLH-2 expression construct

A 1.5 Kb genomic upstream sequence of the HLH-2 [KY21.Chr4.865 (*85*)] gene was amplified using the primers 5’-CCTGCAGGTCGACTCTAGAGAATGTGAACGCTCCAGATG*-3’* and 5’-TCGGAGGAAGCCATTGGTACACTCATGTTGATAGAAGTTTGTATAAAAC*-3’*. NEBuilder HiFi Assembly (NEB, Ipswich, Mass.) was used to join the promoter region and the first 4 amino acids in frame to a *Ciona* optimized RFP vector (pSP72-1.27 mRFPci) (*86*).

### In situ hybridization

To label mRNA fluorescently in whole mount *Ciona* larvae or embryos, the hybridization chain reaction (HCR) in situ method (Molecular Instruments) was followed, as previously described (*13*). Genes for which probe sets were designed and their sequence identifiers are as follows: vesicular acetylcholine transporter (VACHT; NM_001032789.1), motor neuron homeobox (MNX; KH2012:KH.L128.12) lim-domain homeobox 1/5 (LHX/5; KH2012.L107.7), vesicular GABA transporter (VGAT; NM_001032573.1), and Orthopedia (OTP; XM_009862516.3).

### Immunolabeling

Labeling with antibodies was performed as previously described (*5*). Primary antibodies used were as follows: mCherry (rat; life Technologies), Kaede (rabbit; MBL), and GFP (mouse; Invitrogen). Appropriate fluorescent Alexa Fluor secondary antibodies were purchased from Invitrogen.

### SU5402 treatment

SU5402 (Sigma) was diluted in DMSO to make a stock solution at 10µg/µL, which was diluted to a working solution ranging from 20-40µM. DMSO-only served as a negative control. The drug was added to seawater-filled dishes containing late-neurula stage embryos, which had resulted from a cross between wild type eggs and stable transgenic VGAT>Kaede sperm. Thirty minutes after addition of the drug, the embryos were washed four times with filtered seawater, then allowed to reach the larval stage, when they were fixed for immunolabeling and then prepared for confocal imaging.

### Ciona connectome

Analysis of *Ciona* connectomic data and generation of connectivity diagrams were performed using Cytoscape (*87*). The connectivity matrix is available for download from (*3*). Reconstruction of *Ciona* CNS domains using serial section EM images of the connectome project were generated using the Cionectome Viewer on Morphonet (https://morphonet.org/).

### Phylogenetic analysis

Protein-protein BLAST was performed using human Ptf1a (NP_835455.1) in the NCBI non-redundant protein sequences database. Eighteen proteins in 9 diverse chordate taxa were selected for phylogenetic analysis (mouse ptf1a: NP_061279.2, chicken ptf1a: XP_015137641.1, frog ptf1a: NP_001167491.1, hagfish ptf1a: BCS79892.1, lancelet ptf1a: CAH1239013.1, lamprey ptf1a: XP_032824760.1, human fer3-like: EAW93713.1, mouse fer3-like: NP_277057.1, chicken fer3-like: XP_040520525.1, frog fer3-like: XP_018122450.1, *Ciona* ptf1a-related: XP_026693718.1, *Styela* fer3-like: XP_039261045.1, human twist-related: CAA67664.1, mouse twist-related: NP_035788.1, frog twist-related: NP_001079352.1, lamprey twist-related: XP_032807567.1).

The evolutionary history was inferred by using the Maximum Likelihood method and Jones-Taylor-Thornton (JTT) matrix-based model with 500 bootstrap replicates (*88*). Initial tree for the heuristic search was obtained automatically by applying Neighbor-Join and BioNJ algorithms to a matrix of pairwise distances estimated using the JTT model, and then selecting the topology with superior log likelihood value. Evolutionary analyses were conducted in MEGA11 (*89*).

### Transgenesis by electroporation

Plasmids [ASIC> and NSCL-2>GFP reporter constructs, and VGAT>jGCaMP6f (*90*)] were electroporated into 1-cell stage embryos, as described (*91*). Briefly, fertilized eggs were first dechorinated with 1% sodium thioglycolate and 0.05% protease E, and then washed extensively with seawater. The dechorionated fertilized eggs in 300 𝛍L of seawater were mixed with 50 𝛍g DNA in 480 𝛍L 0.77 M mannitol solution and electroporated at 0.05 kV. Fertilized eggs were then washed extensively with sea water and cultured at 18°C in seawater with 0.1% methyl cellulose.

### GCaMP imaging

At the larval stage, transgenics expressing for VGAT>jGCaMP6f were mounted under coverslips raised with ∼80 𝜇m spacers on glass slides in filtered seawater at 24-26 hour post fertilization, as described previously (*12*). GCaMP6f fluorescence was recorded with a Leica DM6B fluorescence microscope using a 20X lens at a frame rate of 10-15 frames per second. The resulting movies were analyzed for temporal changes in fluorescence in selected regions of interest corresponding to the AMG cells using ImageJ (*92*). Normalized fluorescence values (ΔF/F_0_) were calculated as previously described (*12*).

### Single-cell RNA sequencing analysis

The raw sequencing reads of two previously published larval datasets (SRP198321 and SRP166883) were retrieved from Sequence Read Archive (*30*, *31*). The reads were mapped to the KY2021 reference genome using CellRanger 8.0.1 (*93*, *94*). The Datasets were integrated with a linearly-decoded variational auto-encoder (LDVAE) (*95*). The optimal configuration of the hyperparameters of the LDVAE were chosen by sweeping the hyperparameter space. Leiden clustering was performed using ScanPy (*96*). Differential gene expression analysis was performed using scvi-tool 1.2 (*97*). The name of the closest human BLAST result (e-value < 0.05) from the UniProt database was used to annotate the KY21 genes.

### Cross-species analysis

One-to-one protein orthology between the mouse M10 genome (GRCm38.p4) and the *Ciona* KY2021 genome was identified using OrthoFinder (*98*). The mouse cerebellum datasets were mapped to the KY2021 genome using ortholog genes as previously described (*68*, *99*). The transformed mouse datasets were integrated with the *Ciona* dataset generated in the previous section using scvi-tool 1.2 and ScanPy. Mean dot products between the expression vector in the LDVAE latent space were calculated for every possible pair of *Ciona* and mouse clusters.

## Acknowledgements

We thank Anna Di Gregorio for the ASIC plasmid, and Alberto Stolfi for the Hox10 plasmid. We thank Yasunori Sasakura and the CITRES team for providing the stable VGAT>Kaede line. We also thank Ian Meinertzhagen for his helpful comments on the manuscript.

## Data and materials availability

The integrated *Ciona robusta* larva stage single cell RNA sequencing dataset and the complete differential expression data for all clusters shown in Figure 4C are deposited at Zenodo (https://doi.org/10.5281/zenodo.15320250).

