## Supplemental Data for "The dorsal/ventral subdivision of the hindbrain predates the tunicate/vertebrate split"

**Movie S1. pATENs are mechanosensitive**. Three *Ciona* larvae responding to being touched at pATENs with a fine probe. The movie plays in real time.

**Movie S2. Spiking activity of AMGs**. Calcium transients in a *Ciona* larva expressing VGAT>jGCaMP6f. The movie shows 47 seconds of recording at 8.7 frames/second. The movie plays at 5X speed.

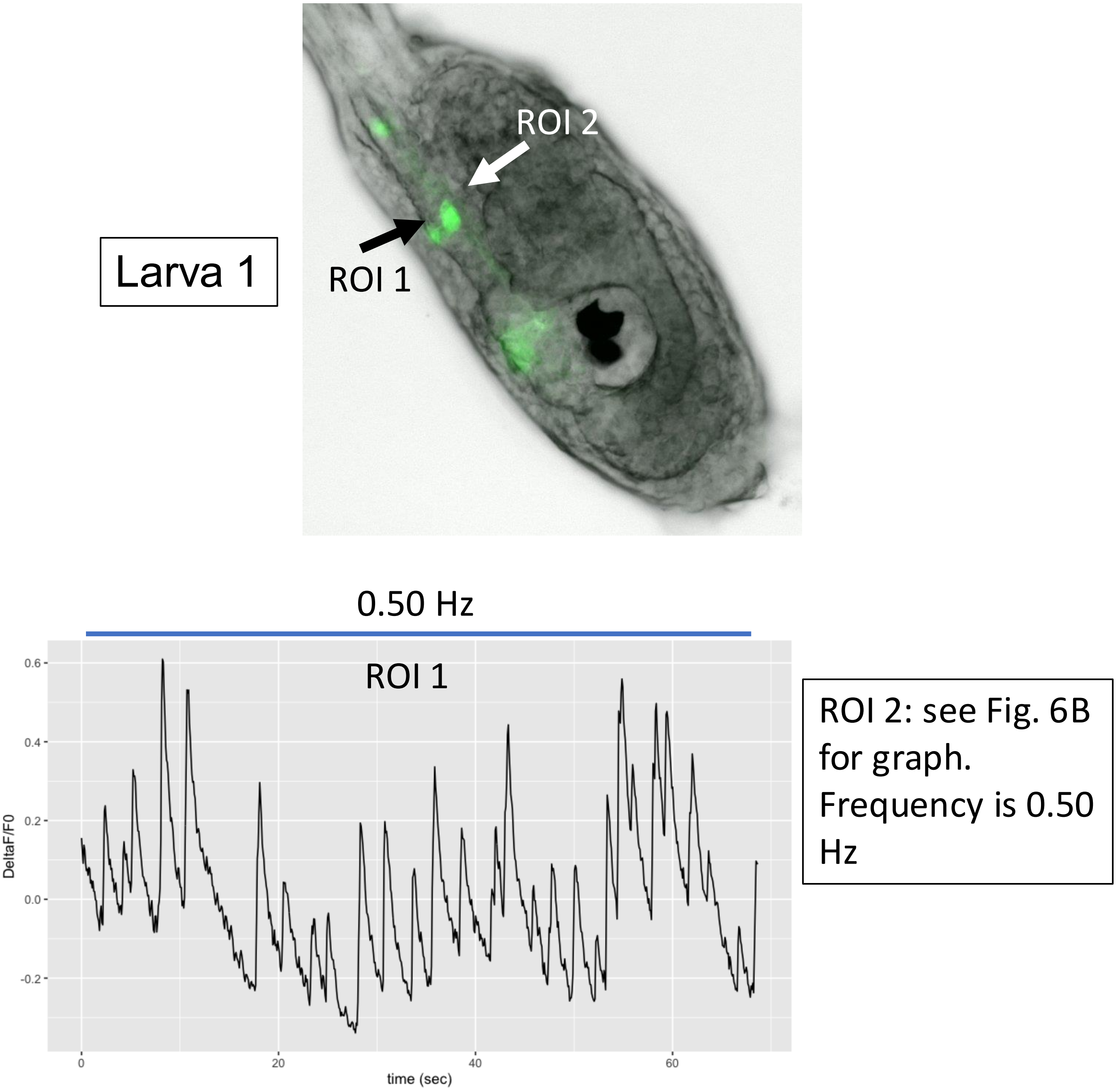

**Fig S1. Spontaneous spiking activity in the inhibitory AMG neurons.** Data from five larvae transfected with the plasmid VGAT>GCaMP6f. Arrows in left panels indicate the neuron(s) analyzed. Right panels show plots of normalized GCaMP6f fluorescence ($\Delta F/F_0$). Two regions of interest (ROI), corresponding to separate neurons, were analyzed for larva 1. In all other larvae, a single ROI was analyzed (arrows).

Larva 2

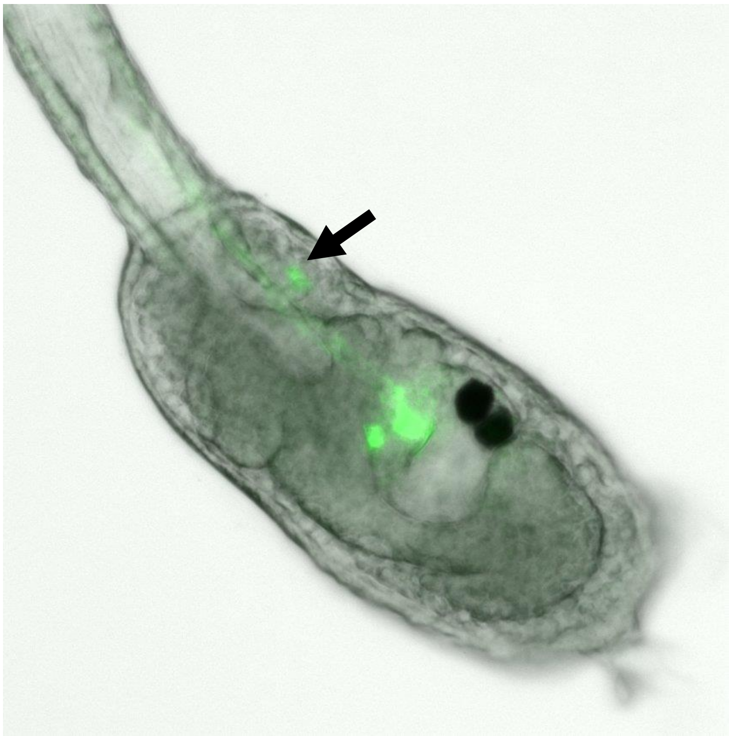

0.39 Hz

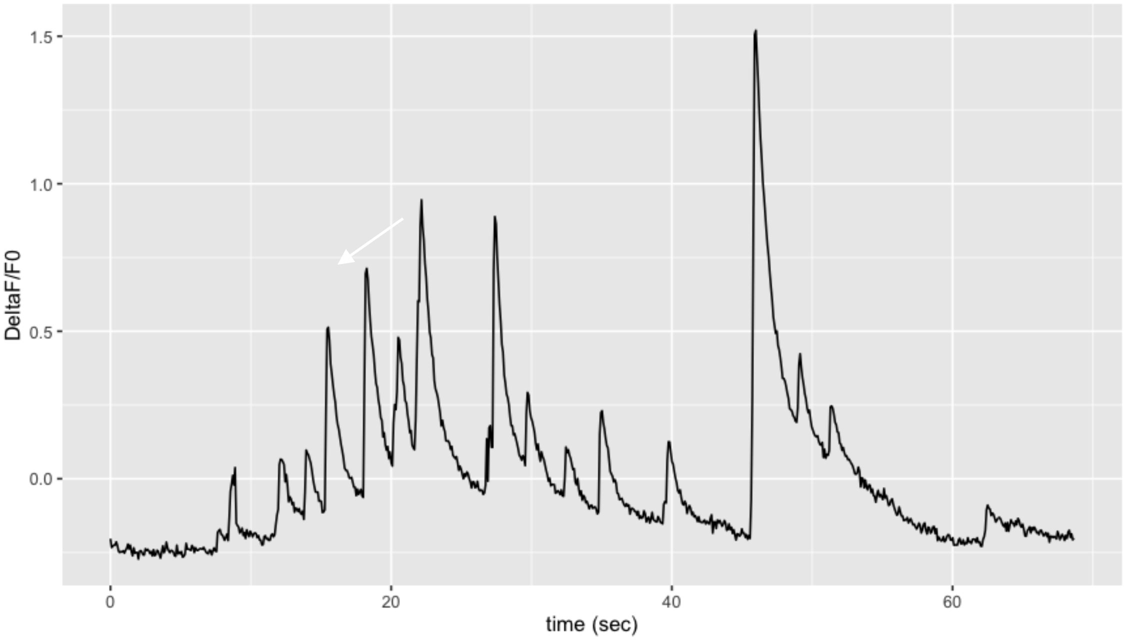

**Fig S1 (cont.)**

Larva 3

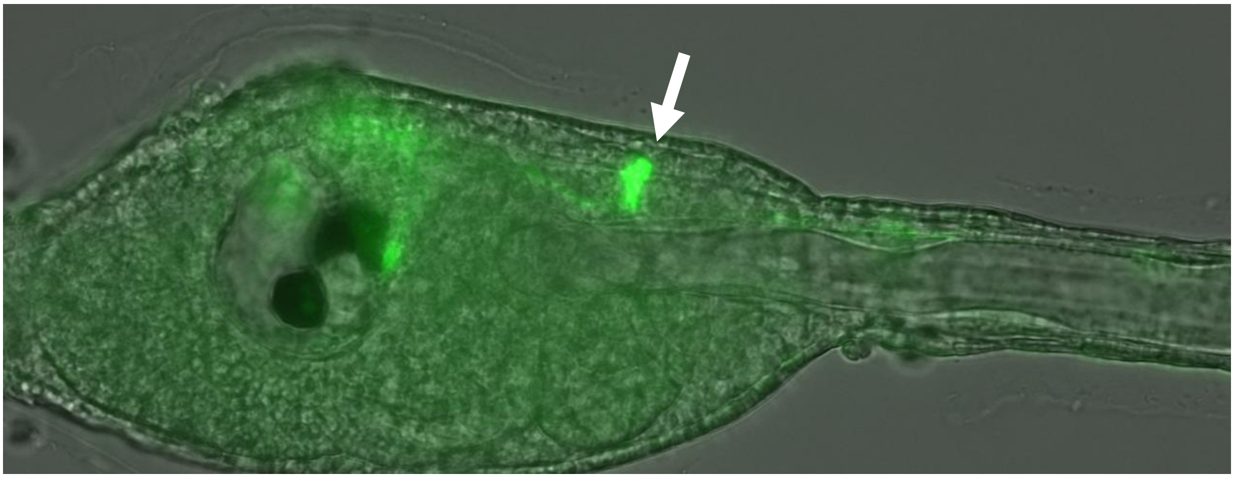

0.53 Hz

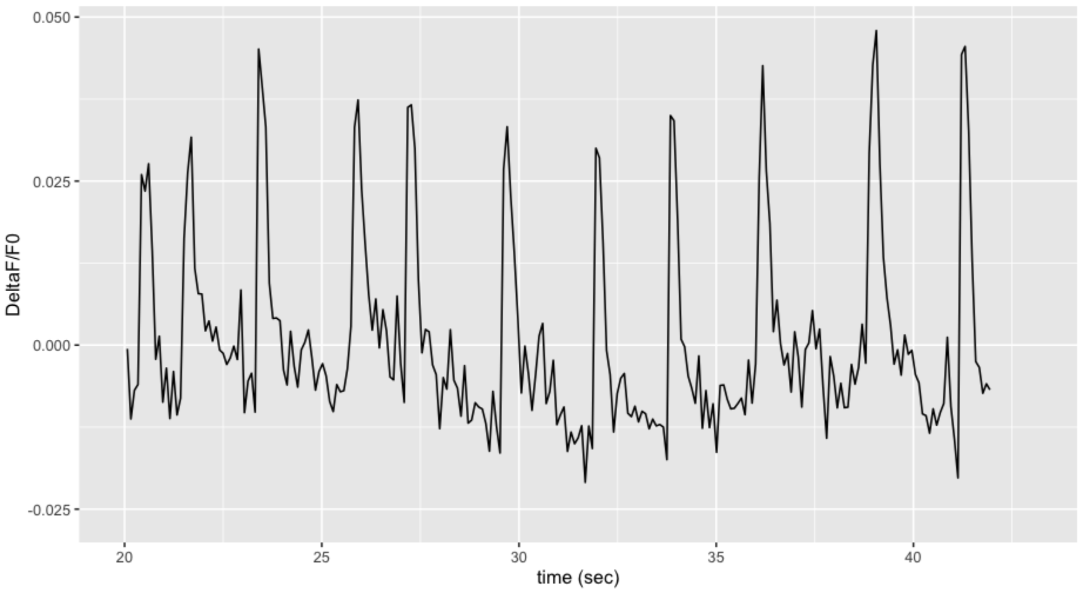

**Fig S1 (cont.)**

Larva 4

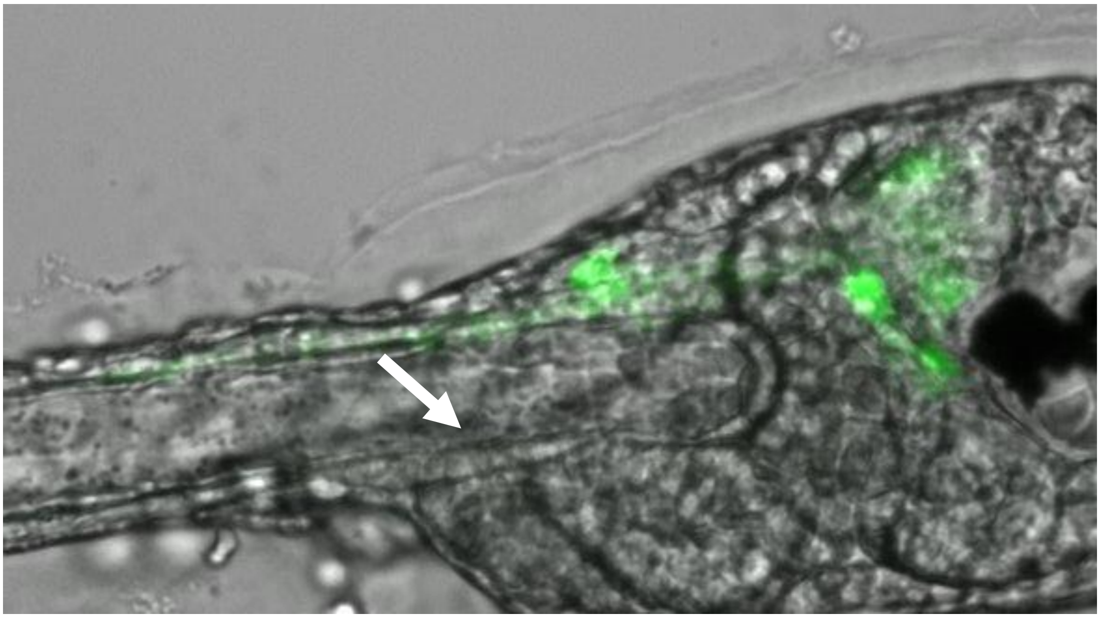

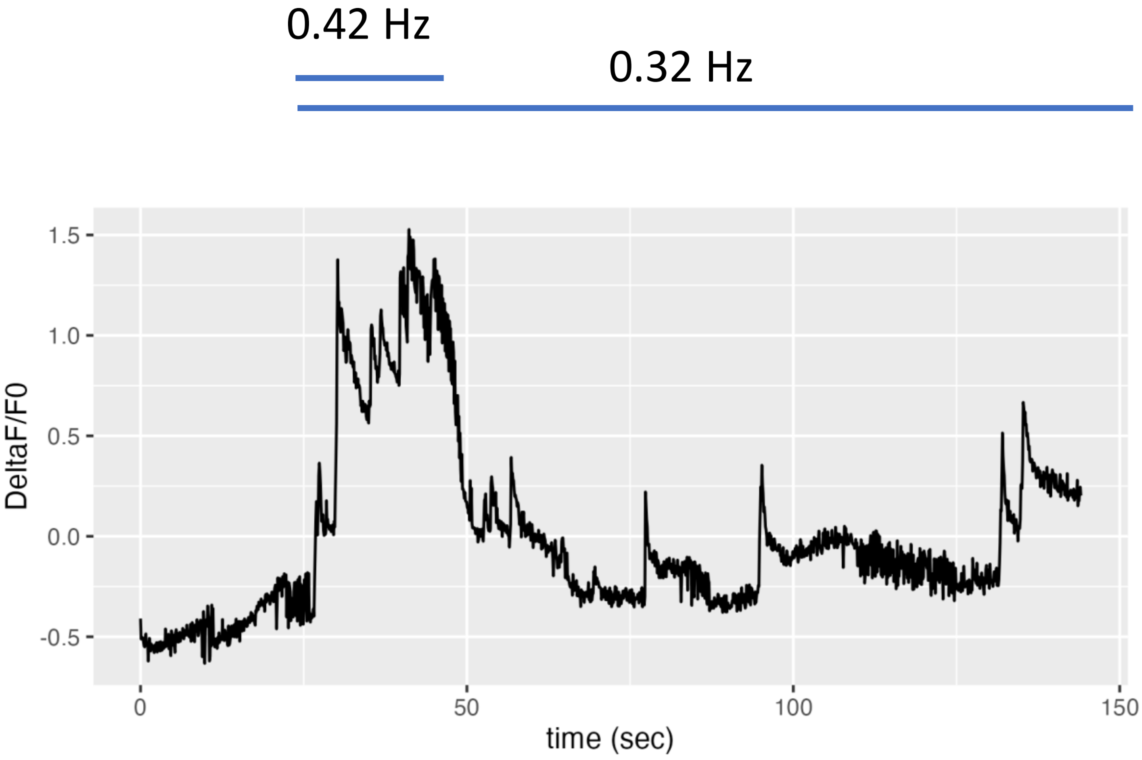

**Fig S1 (cont.)**

Larva 5

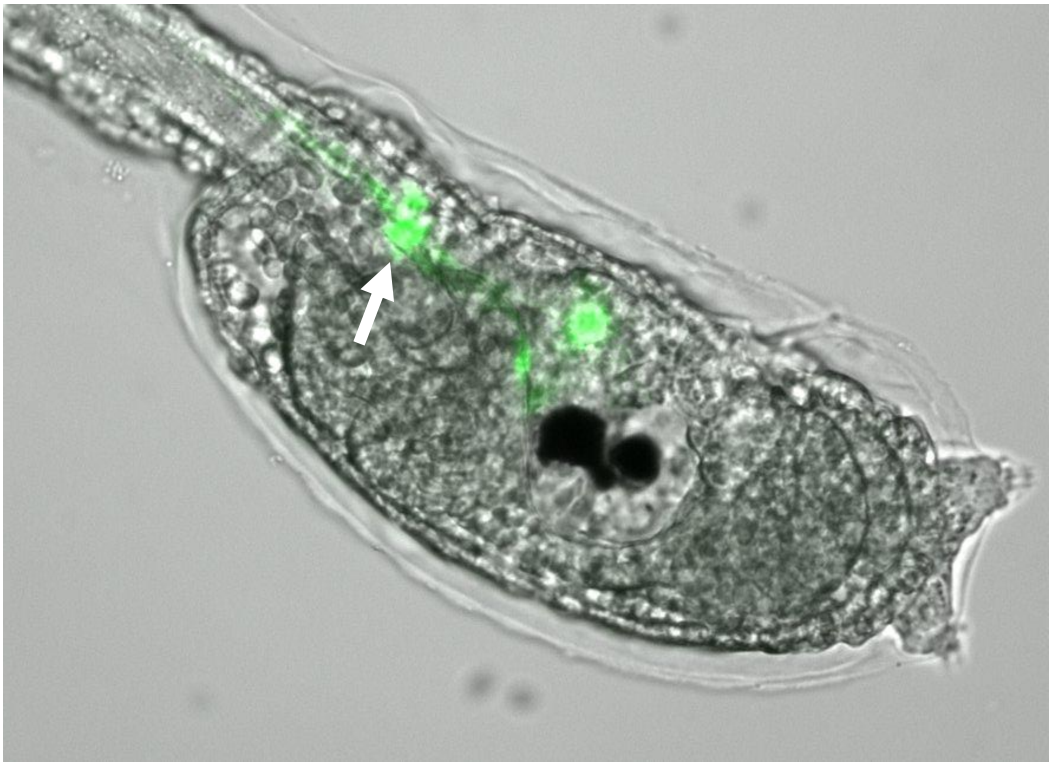

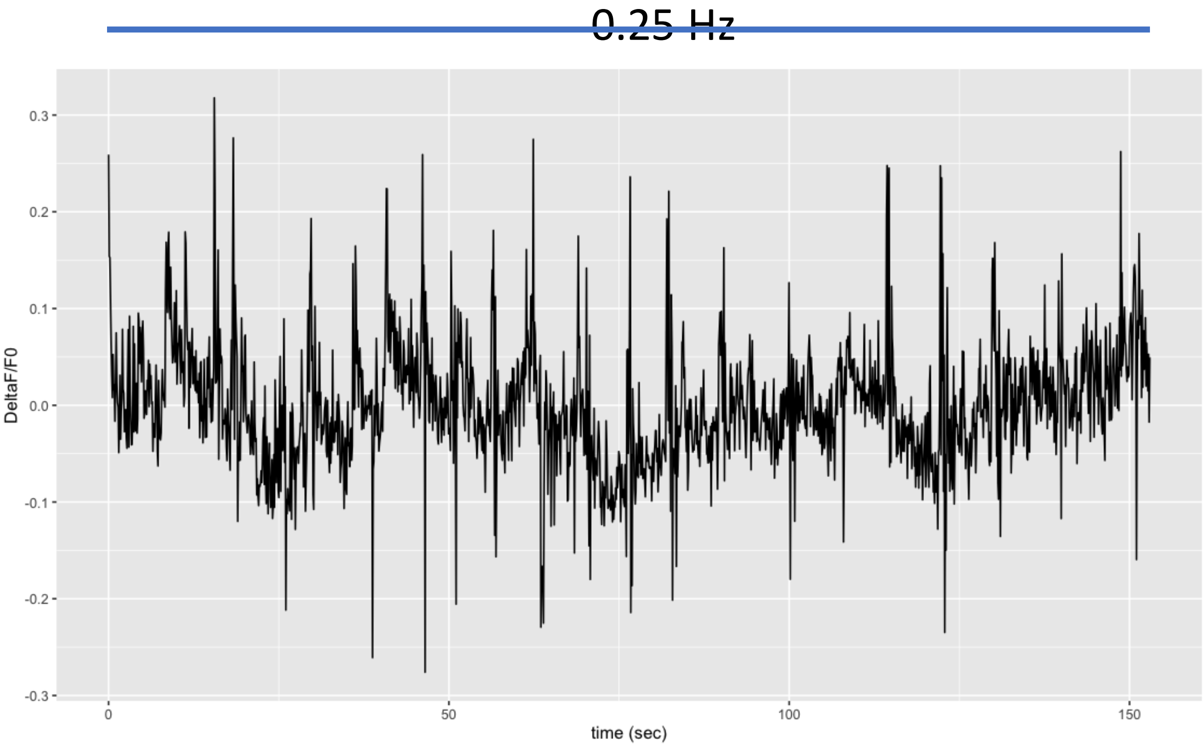

**Fig S1 (cont.)**

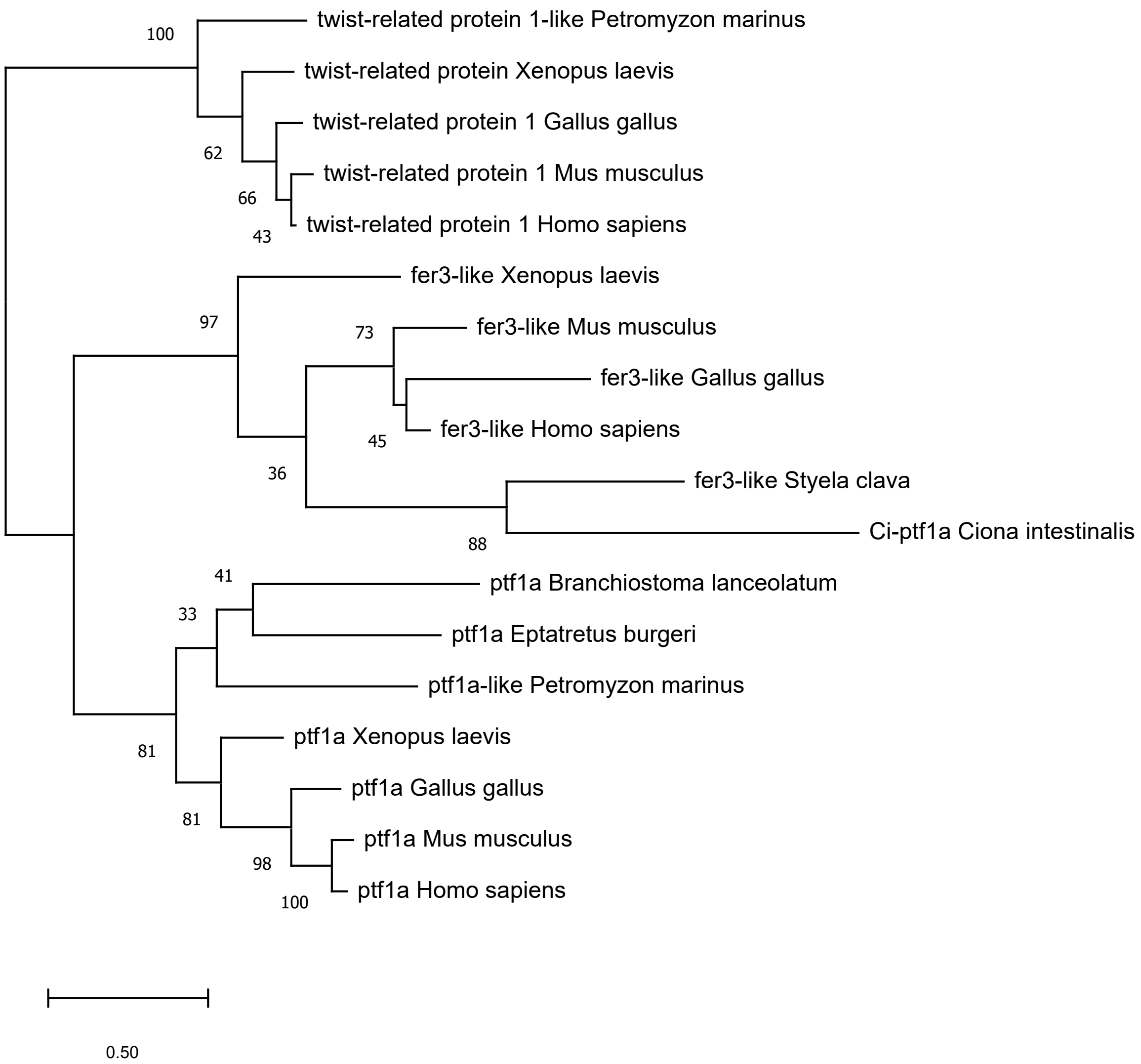

**Fig S2.** Phylogenic analysis of chordate Ptf1a family members. The putative *Ciona* Ptf1a ortholog (yellow highlight) groups with the fer-like class of transcription factors. Despite the absence of a Ptf1a ortholog in Ciona, an orthologous gene is present in cephalochordates (Branchiostoma lanceolatum; blue highlight).

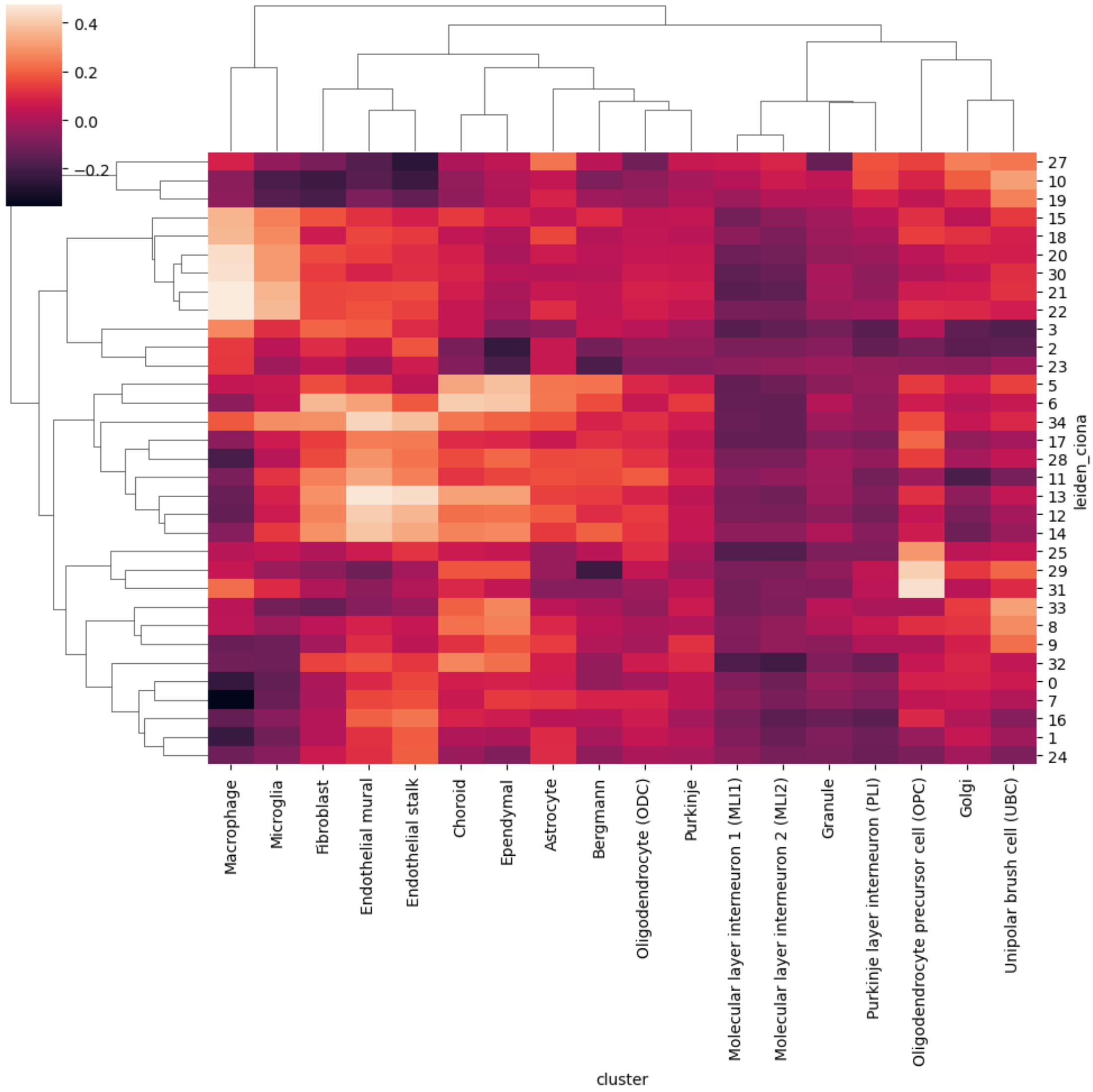

**Fig S3.** Heat map of mean cosine similarity between annotated mouse cerebellum single cell RNA sequencing clusters and *Ciona* clusters. Several *Ciona* clusters showed considerable expression similarities with mouse cerebellar clusters. However, mouse Purkinje cells cluster does not show an obvious homologous clusters in *Ciona*.

| TABLE 1 | | | | |
|---|---|---|---|---|
| Gene ID (KY2021) | lfc_mean | Best Hit Human: uniprot | Best Hit Human: fullname | Notes on expression of vertebrate best hit |
| KY21.Chr1.2051 | 7.043459 | P78348 | Acid-sensing ion channel 1 {ECO:0000303\|PubMed:10798398} | Enriched in Purkinje Cells; doi:10.1073/pnas.94.4.1459; doi:10.1038/386173a0 |
| KY21.Chr7.124 | 6.608743 | Q8TBE7 | Solute carrier family 35 member G2{ECO:0000312\|HGNC:HGNC:28480} | |
| KY21.Chr1.1244 | 6.595257 | O95672 | Endothelin-converting enzyme-like 1 | Stongly expressed in Purkinje cells, but not elsewhere in cellebellum; doi: 10.1016/s0306-4522(98)00692-7 |
| KY21.Chr4.865 | 6.56292 | Q02577 | Helix-loop-helix protein 2 (HLH-2) | Expressed in developing Purkinje cells; doi: 10.1016/0169-328x(95)00282-w |
| KY21.Chr2.793 | 6.445879 | Q9H598 | Vesicular inhibitory amino acid transporter {ECO:0000303\|PubMed:12031963} | Widely expressed in CNS inlcuding Purkinje cells; doi: 10.3389/fncel.2013.00286 |
| KY21.Chr11.1031 | 6.37433 | O95076 | Homeobox protein aristaless-like 3 | |
| KY21.Chr2.267 | 6.364623 | Q9HB71 | Calcyclin-binding protein | highly enriched in Purkinje cells; DOI: 10.1177/002215540004800903 |
| KY21.Chr1.2142 | 6.361967 | Q99680 | G-protein coupled receptor 22 | |
| KY21.Chr11.1275 | 6.349948 | Q9H2C1 | LIM/homeobox protein Lhx5 | Expressed in Purkinje cells:doi.org/10.1007/s00418-023-02251-z |
| KY21.Chr4.611 | 6.336725 | NA | NA | |
| KY21.Chr6.570 | 6.292621 | NA | NA | |
| KY21.Chr10.677 | 6.257514 | NA | NA | |
| KY21.Chr11.1122 | 6.237363 | P33032 | Melanocortin receptor 5 | |

**table S1. Differentially expressed genes in the inhibitory AMG single-cell transcriptome cluster.**

Differential expression (DE) is listed in descending order of $\log_2$ mean fold-change (*mean lfc*). DE genes with a *mean lfc* less than 4.8 and an absolute expression level (*raw normalized mean*) less than 0.2 were excluded. Compiled *mean lfc* for all annotated genes in all clusters is available at https://doi.org/10.5281/zenodo.15320250. Genes with no apparent vertebrate homolog as assessed by BLAST analysis (cutoff E< 0.05) are highlighted in orange. Yellow highlighting corresponds to genes that were used to identify the cluster (VGAT, Lhx1/5 and Otp). Select genes with known expression and/or function in the cerebellum are highlighted in green, along with a reference. Translated *Ciona* gene models [KY2021} can be downloaded from http://ghost.zool.kyoto-u.ac.jp/datas/HT.KY21Gene.protein.2.fasta.zip.

| | | | | |
|---|---|---|---|---|
| KY21.Chr12.770 | 6.168701 | Q8N475 | Follistatin-related protein 5 | Highly expressed in cerebellum; doi.org/10.1111/cga.12022 |
| KY21.Chr3.488 | 6.12561 | NA | NA | |
| KY21.Chr3.1549 | 6.117251 | P21266 | Glutathione S-transferase Mu 3 | |
| KY21.Chr14.678 | 6.102291 | Q96EP9 | Sodium/bile acid cotransporter 4 | |
| KY21.Chr12.1019 | 6.091909 | Q5TCZ1 | SH3 and PX domain-containing protein 2A | |
| KY21.Chr1.783 | 6.048831 | P28329 | Choline O-acetyltransferase | |
| KY21.Chr11.1143 | 6.032415 | P32297 | Neuronal acetylcholine receptor subunit alpha-3 | |
| KY21.Chr2.721 | 6.024844 | Q13394 | Putative nucleotidy ltransferase MAB21L1{ECO:0000305} | |
| KY21.Chr10.378 | 6.002219 | Q05901 | Neuronal acetylcholine receptor subunit beta-3 | |
| KY21.Chr3.486 | 5.991252 | NA | NA | |
| KY21.Chr1.1224 | 5.978268 | P23352 | Anosmin-1 {ECO:0000303\|PubMed:8832397, ECO:0000312\|HGNC:HGNC:6211} | Stimulates outgrowth and branching of developing Purkinje axons; doi:10.1016/j.neuroscience.2008.10.022 |
| KY21.Chr4.240 | 5.955939 | NA | NA | |
| KY21.Chr4.1089 | 5.944547 | Q9NP94 | Zinc transporter ZIP2 | |
| KY21.Chr1.566 | 5.940061 | A7MD48 | Serine/arginine repetitive matrix protein 4 | |
| KY21.Chr7.1029 | 5.936464 | Q8TBB6 | Solute carrier family 7 member 14 | |
| KY21.Chr12.180 | 5.926925 | P16870 | Carboxypeptidase E | |
| KY21.Chr3.1047 | 5.895681 | P30988 | Calcitonin receptor {ECO:0000305} | Enriched in developing cerebellum; https://doi.org/10.1002/cne.10478 |
| KY21.Chr4.864 | 5.879049 | P36383 | Gap junction gamma-1 protein | |
| KY21.Chr10.367 | 5.876298 | P16519 | Neuroendocrine convertase 2 | |

**table S1, cont.**

| | | | |
|---|---|---|---|
| KY21.Chr5.104 | 5.84712 | A6NJT0 | Homeobox protein unc-4 homolog |
| KY21.Chr2.1203 | 5.827642 | P21579 | Synaptotagmin-1 {ECO:0000303\|PubMed:25705886} |
| KY21.Chr14.230 | 5.807533 | Q6UXK2 | Immunoglobulin superfamily containing leucine-rich repeat protein 2 |
| KY21.Chr3.590 | 5.802678 | NA | NA |
| KY21.Chr3.20 | 5.799918 | A6NKL6 | Transmembrane protein 200C |
| KY21.Chr6.500 | 5.796391 | Q5TAB7 | Protein ripply2 |
| KY21.Chr5.1117 | 5.786946 | Q13018 | Secretory phospholipase A2 receptor |
| KY21.Chr1.1 | 5.761394 | Q92913 | Fibroblast growth factor 13 {ECO:0000305} |
| KY21.Chr1.1910 | 5.744116 | P24046 | Gamma-aminobutyric acid receptor subunit rho-1 {ECO:0000303\|PubMed:1849271} |
| KY21.Chr9.601 | 5.730854 | Q8N608 | Inactive dipeptidyl peptidase 10 |
| KY21.Chr2.491 | 5.725424 | NA | NA |
| KY21.Chr6.24 | 5.721511 | P32297 | Neuronal acetylcholine receptor subunit alpha-3 |
| KY21.Chr3.1580 | 5.709742 | NA | NA |
| KY21.Chr10.463 | 5.709182 | O95471 | Claudin-7 |
| KY21.Chr14.449 | 5.69499 | NA | NA |
| KY21.Chr2.1365 | 5.659719 | Q92930 | Ras-related protein Rab-8B |
| KY21.Chr3.9 | 5.642222 | Q96FS4 | Signal-induced proliferation-associated protein 1 |
| KY21.Chr5.81 | 5.641432 | Q9UNE2 | Rab effector Noc2 |
| KY21.Chr7.238 | 5.633389 | P36383 | Gap junction gamma-1 protein |
| KY21.Chr1.1844 | 5.633156 | Q9Y4B5 | Microtubule cross-linking factor 1 |
| KY21.Chr5.603 | 5.623641 | O14994 | Synapsin-3 |
| KY21.Chr10.618 | 5.61895 | NA | NA |

**table S1, cont.**

| | | | |
|---|---|---|---|
| KY21.Chr11.123 | 5.602503 | Q9UI40 | Sodium/potassium/calcium exchanger 2 |
| KY21.Chr14.382 | 5.602118 | O00591 | Gamma-aminobutyric acid receptor subunit pi {ECO:0000250\|UniProtKB:O09028} |
| KY21.Chr1.217 | 5.600185 | Q6IQ22 | Ras-related protein Rab-12 |
| KY21.Chr3.580 | 5.592871 | Q9ULB1 | Neurexin-1 |
| KY21.Chr3.1466 | 5.569221 | P05771 | Protein kinase C beta type |
| KY21.Chr8.836 | 5.566675 | NA | NA |
| KY21.Chr11.1061 | 5.549929 | Q69YW2 | Protein stum homolog |
| KY21.Chr11.1179 | 5.541953 | NA | NA |
| KY21.Chr12.830 | 5.540315 | NA | NA |
| KY21.Chr3.886 | 5.537454 | O76038 | Secretagogin |
| KY21.Chr7.1153 | 5.529147 | NA | NA |
| KY21.Chr1.422 | 5.523847 | Q9H4W6 | Transcription factor COE3 |
| KY21.Chr11.193 | 5.521917 | Q6ZNA5 | Ferric-chelate reductase 1 |
| KY21.Chr3.1637 | 5.5129 | Q15907 | Ras-related protein Rab-11B |
| KY21.Chr3.667 | 5.507032 | Q99622 | Protein C10 |
| KY21.Chr13.97 | 5.494586 | P48544 | G protein-activated inward rectifier potassium channel 4 |
| KY21.Chr1.1592 | 5.492125 | O75325 | Leucine-rich repeat neuronal protein 2 |
| KY21.Chr8.1318 | 5.479926 | A6NHT5 | Homeobox protein HMX3 |
| KY21.Chr8.468 | 5.474479 | Q6PUV4 | Complexin-2 |
| KY21.Chr9.574 | 5.43528 | P28472 | Gamma-aminobutyric acid receptor subunit beta-3 {ECO:0000303\|PubMed:8382702} |
| KY21.Chr1.1459 | 5.419416 | Q9HCR9 | Dual 3',5'-cyclic-AMP and -GMP phosphodiesterase 11A |
| KY21.Chr9.761 | 5.414707 | Q12879 | Glutamate receptor ionotropic, NMDA 2A {ECO:0000305} |
| KY21.Chr2.1142 | 5.404658 | Q16555 | Dihydropyrimidinase-related protein 2 |
| KY21.Chr6.262 | 5.387043 | Q9NSD7 | Relaxin-3 receptor 1 |

**table S1, cont.**

| | | | |
|---|---|---|---|
| KY21.Chr6.231 | 5.369979 | O95948 | One cut domain family member 2 |
| KY21.Chr12.916 | 5.364193 | Q13641 | Trophoblast glycoprotein |
| KY21.Chr2.1366 | 5.361039 | P51787 | Potassium voltage-gated channel subfamily KQT member 1 {ECO:0000305} |
| KY21.Chr11.360 | 5.35556 | NA | NA |
| KY21.Chr1.191 | 5.333432 | P62166 | Neuronal calcium sensor 1 |
| KY21.Chr7.624 | 5.322394 | NA | NA |
| KY21.Chr5.876 | 5.30879 | P59768 | Guanine nucleotide-binding protein G(I)/G(S)/G(O) subunit gamma-2 |
| KY21.Chr1.2010 | 5.303323 | Q09470 | Potassium voltage-gated channel subfamily A member 1 {ECO:0000305} |
| KY21.Chr3.1172 | 5.27283 | Q8NDX2 | Vesicular glutamate transporter 3 {ECO:0000303\|PubMed:12151341} |
| KY21.Chr11.997 | 5.25814 | P50238 | Cysteine-rich protein 1 |
| KY21.Chr2.125 | 5.257552 | O43581 | Synaptotagmin-7 {ECO:0000305} |
| KY21.Chr12.1064 | 5.257227 | P21452 | Substance-K receptor |
| KY21.Chr7.1221 | 5.247487 | Q99259 | Glutamate decarboxylase 1 |
| KY21.Chr2.733 | 5.238503 | NA | NA |
| KY21.Chr7.252 | 5.235325 | P14416 | D(2) dopamine receptor |
| KY21.Chr8.957 | 5.211644 | Q9UBP4 | Dickkopf-related protein 3 |
| KY21.Chr10.147 | 5.201596 | O15394 | Neural cell adhesion molecule 2 |
| KY21.Chr7.1003 | 5.18675 | NA | NA |
| KY21.Chr1.1311 | 5.171793 | Q9NNX6 | CD209 antigen |
| KY21.Chr1.1090 | 5.169981 | NA | NA |
| KY21.Chr3.140 | 5.15821 | P0C851 | Phosphoinositide-interacting protein |
| KY21.Chr4.1039 | 5.15735 | Q9H2E6 | Semaphorin-6A |
| KY21.Chr12.637 | 5.147467 | P0DP23 | Calmodulin-1 {ECO:0000312\|HGNC:HGNC:1442} |

table S1, cont.

| | | | | |
|---|---|---|---|---|
| KY21.Chr10.515 | 5.144052 | Q9Y3Q4 | Potassium/sodium hyperpolarization-activated cyclic nucleotide-gated channel 4 | |
| KY21.Chr9.379 | 5.134594 | Q13237 | cGMP-dependent protein kinase 2 | |
| KY21.Chr11.1113 | 5.131234 | P11229 | Muscarinic acetylcholine receptor M1 | |
| KY21.Chr4.922 | 5.11491 | Q9UL51 | Potassium/sodium hyperpolarization-activated cyclic nucleotide-gated channel 2 | |
| KY21.Chr9.470 | 5.113177 | Q9UBS5 | Gamma-aminobutyric acid type B receptor subunit 1 | |
| KY21.Chr9.1108 | 5.110392 | P37288 | Vasopressin V1a receptor | |
| KY21.Chr11.804 | 5.109746 | Q96SJ8 | Tetraspanin-18 {ECO:0000305} | |
| KY21.Chr8.227 | 5.109483 | Q9BZV3 | Interphotoreceptor matrix proteoglycan 2 | |
| KY21.Chr5.741 | 5.107118 | NA | NA | |
| KY21.Chr5.217 | 5.096029 | P35372 | Mu-type opioid receptor | |
| KY21.Chr2.686 | 5.08231 | Q02153 | Guanylate cyclase soluble subunit beta-1 {ECO:0000305} | |
| KY21.Chr5.906 | 5.073726 | Q9NYY8 | FAST kinase domain-containing protein 2, mitochondrial | |
| KY21.Chr14.175 | 5.06331 | Q9UHG0 | Doublecortin domain-containing protein 2 | |
| KY21.Chr14.647 | 5.053997 | Q5XKR4 | Homeobox protein orthopedia | Expressed in inhibitory cerebellar cells:doi.org/10.1007/s00418-023-02251-z |
| KY21.Chr1.863 | 5.053265 | P0DP23 | Calmodulin-1 {ECO:0000312\|HGNC:HGNC:1442} | |
| KY21.Chr1.964 | 5.045961 | Q9H4D0 | Calsyntenin-2 {ECO:0000303\|PubMed:12498782} | |

**table S1, cont.**

| | | | |
|---|---|---|---|
| KY21.Chr10.1007 | 5.033205 | Q9UMX3 | Bcl-2-related ovarian killer protein {ECO:0000303\|PubMed:11034351} |
| KY21.Chr11.890 | 5.014723 | Q9BTD3 | Transmembrane protein 121 |
| KY21.Chr13.132 | 5.01279 | P48066 | Sodium- and chloride-dependent GABA transporter 3 |
| KY21.Chr13.457 | 5.012593 | Q9UBR4 | LIM/homeobox protein Lhx3 |
| KY21.Chr2.1082 | 4.981819 | P07101 | Tyrosine 3-monooxygenase |
| KY21.Chr14.837 | 4.977802 | Q9HCJ2 | Leucine-rich repeat-containing protein 4C |
| KY21.Chr14.109 | 4.969039 | Q96QF0 | Rab-3A-interacting protein |
| KY21.Chr8.1088 | 4.967403 | P0DPH7 | Tubulin alpha-3C chain |
| KY21.Chr1.1208 | 4.963042 | P35372 | Mu-type opioid receptor |
| KY21.Chr13.308 | 4.950118 | NA | NA |
| KY21.Chr3.561 | 4.933967 | Q7Z553 | MAM domain-containing glycosylphosphatidylinositol anchor protein 2 |
| KY21.Chr11.400 | 4.932946 | Q9BZE3 | BarH-like 1 homeobox protein |
| KY21.Chr4.127 | 4.93143 | Q8N2C7 | Protein unc-80 homolog |
| KY21.Chr9.1072 | 4.927906 | Q9Y6C2 | EMILIN-1 |
| KY21.Chr8.581 | 4.921591 | P49221 | Protein-glutamine gamma-glutamyltransferase 4 |
| KY21.Chr3.263 | 4.905591 | O15344 | E3 ubiquitin-protein ligase Midline-1 |
| KY21.Chr12.1099 | 4.894771 | P0DP23 | Calmodulin-1 {ECO:0000312\|HGNC:HGNC:1442} |
| KY21.Chr4.818 | 4.874436 | Q9ULZ9 | Matrix metalloproteinase-17 |
| KY21.Chr9.1175 | 4.863217 | Q9NQX5 | Neural proliferation differentiation and control protein 1 |
| KY21.Chr9.371 | 4.862268 | Q9NSD5 | Sodium- and chloride-dependent GABA transporter 2 |

**table S1, cont.**

| | | | |
|---|---|---|---|
| KY21.Chr4.553 | 4.861459 | Q16566 | Calcium/calmodulin-dependent protein kinase type IV |
| KY21.Chr1.2267 | 4.858945 | NA | NA |
| KY21.Chr9.941 | 4.8581 | Q86SS6 | Synaptotagmin-9 |
| KY21.Chr1.1533 | 4.850421 | P49795 | Regulator of G-protein signaling 19 |
| KY21.Chr2.768 | 4.835589 | NA | NA |
| KY21.Chr6.372 | 4.81905 | NA | NA |
| KY21.Chr2.847 | 4.811907 | NA | NA |
| KY21.Chr1.291 | 4.782984 | P0DP23 | Calmodulin-1 {ECO:0000312\|HGNC:HGNC:1442} |
| KY21.Chr2.734 | 4.778576 | P42263 | Glutamate receptor 3 {ECO:0000305} |
| KY21.Chr2.1462 | 4.746126 | Q9NPC2 | Potassium channel subfamily K member 9 |
| KY21.Chr3.1525 | 4.715206 | P0DP23 | Calmodulin-1 {ECO:0000312\|HGNC:HGNC:1442} |
| KY21.Chr1.1655 | 4.714397 | Q8WUI4 | Histone deacetylase 7 |
| KY21.Chr2.441 | 4.70557 | Q9BXA6 | Testis-specific serine/threonine-protein kinase 6 |
| KY21.Chr4.522 | 4.695408 | Q8N441 | Fibroblast growth factor receptor-like 1 |
| KY21.Chr6.434 | 4.686903 | Q92886 | Neurogenin-1 |
| KY21.Chr7.1218 | 4.677054 | Q8IZ57 | Neurensin-1 |
| KY21.Chr13.368 | 4.673108 | Q9GZU5 | Nyctalopin |
| KY21.Chr10.944 | 4.662253 | P36383 | Gap junction gamma-1 protein |
| KY21.Chr2.1269 | 4.597596 | Q9UQ13 | Leucine-rich repeat protein SHOC-2 |
| KY21.Chr1.1164 | 4.589523 | NA | NA |
| KY21.Chr2.1327 | 4.580794 | NA | NA |
| KY21.Chr11.476 | 4.566051 | P48544 | G protein-activated inward rectifier potassium channel 4 |
| KY21.Chr1.2192 | 4.556142 | Q9UI40 | Sodium/potassium/calcium exchanger 2 |

**table S1, cont.**

| | | | |
|---|---|---|---|
| KY21.Chr4.918 | 4.554426 | O15068 | Guanine nucleotide exchange factor DBS |
| KY21.Chr1.700 | 4.53225 | NA | NA |
| KY21.Chr5.850 | 4.525064 | Q96A58 | Ras-related and estrogen-regulated growth inhibitor |
| KY21.Chr11.329 | 4.519525 | P30874 | Somatostatin receptor type 2 |
| KY21.Chr9.783 | 4.490294 | Q8N568 | Serine/threonine-protein kinase DCLK2 |
| KY21.Chr13.442 | 4.488406 | Q05586 | Glutamate receptor ionotropic, NMDA 1 {ECO:0000305} |
| KY21.Chr9.1050 | 4.486921 | P26367 | Paired box protein Pax-6 |
| KY21.Chr7.806 | 4.485088 | Q9BRI3 | Proton-coupled zinc antiporter SLC30A2 {ECO:0000305\|PubMed:22733820} |

**table S1, cont.**
